# Redundancy masks functional specificity of SMARCD paralogs in neurodevelopment

**DOI:** 10.64898/2025.12.05.692564

**Authors:** YounJu So, Hadrien Soldati, Laurine Anselme, Simon M. G. Braun

**Affiliations:** Department of Genetic Medicine and Development, Faculty of Medicine, University of Geneva, Switzerland; Institute of Genetics and Genomics in Geneva iGE3, Faculty of Medicine, University of Geneva, Switzerland

## Abstract

The SWI/SNF complex is an essential chromatin remodeler that regulates DNA accessibility during brain development. Through combinatorial assembly of subunits encoded by paralogous genes, SWI/SNF complexes form diverse assemblies across and even within cell types. Although paralogs provide functional redundancy, their mutation in neurodevelopmental disorders suggests specialized roles. Focusing on the core SMARCD subunits, we find that loss of individual paralogs SMARCD1 or SMARCD3 has no impact on cortical development, and even deletion of both paralogs SMARCD1/3 in mice causes only minimal cortical defects, reflecting strong compensatory mechanisms. In neuronal differentiation models, depletion of any combination of paralogs increases the abundance of remaining subunits through protein stabilization rather than transcriptional upregulation. Despite this redundancy, rapid degradation experiments reveal distinct gene regulatory programs for each paralog. In neurons, SMARCD3 uniquely controls oxidative phosphorylation through regulation of metabolic gene networks. Finally, we identify chromatin regulators and transcription factors that associate with SMARCD1- or SMARCD3-containing SWI/SNF complexes, likely conferring paralog-specific targeting. Our findings uncover a dual logic of redundancy and specialization regulating SWI/SNF activity, providing a mechanistic basis for the selective vulnerability of paralog genes in neurodevelopmental disorders.

## Introduction

Neurodevelopmental disorders (NDDs), including autism spectrum disorder (ASD), affect approximately 1-2% of the global population and are characterized by intellectual disability, social deficits, and repetitive behaviors^1^. Large-scale exome sequencing has revealed two major classes of genes recurrently mutated in NDDs: those encoding synaptic proteins and those encoding chromatin regulators^2,3^. While mutations in synaptic genes are expected to affect neuronal activity, the strong enrichment of chromatin regulators is more surprising, as chromatin remodeling is a ubiquitous cellular process. Chromatin, the dynamic structure made of DNA wrapped around nucleosomes, controls genome accessibility and thereby transcriptional regulation^4^. During development, chromatin remodeling orchestrates transcription factor (TF) binding to establish cell type-specific gene expression programs^5^. However, why mutations in broadly expressed chromatin regulators primarily cause brain disorders remains poorly understood^6^.

Among chromatin regulators, genes encoding subunits of the SWI/SNF (also known as BAF) complex are among the most frequently mutated in NDDs^7^. *ARID1B*, a subunit of this complex, is the most recurrently mutated gene in affected individuals^3^. The SWI/SNF complex is an ATP-dependent chromatin remodeler that slides and evicts nucleosomes to activate or repress target genes during development^8^. SWI/SNF complexes are highly conserved from yeast to humans in overall structure and biochemical activity, yet mammalian complexes have undergone extensive expansion in subunit diversity during evolution^9^. Through various combinations of 11 to 13 core subunits encoded by 29 genes, SWI/SNF complexes adopt distinct compositions depending on the cell type and developmental stage. While yeast and flies exhibit only a few different complex assemblies, current estimates suggest that humans can form more than a thousand different SWI/SNF complexes across various tissues^10^. Interestingly, the expansion in genome size and regulatory sequences in mammalian genomes coincides with the expansion in the combinatorial assembly of SWI/SNF subunits. With increased subunit diversity, mammalian SWI/SNF complexes have more varied cell type specific functions during development, enabling unique compositions of the complex to interact with different transcriptional regulators and arrays of chromatin modifications^11^.

Genes such as *ARID1A* and *ARID1B*, which encode alternative versions of the same SWI/SNF subunit, are termed paralogs, having arisen through gene duplication events during evolution. Of the fifteen canonical SWI/SNF subunits, eleven are encoded by multiple paralogous genes, including the ATPase subunits *SMARCA2/4* and the core subunits *SMARCD1/2/3* which share 67-74% identity in their amino acid sequences. This paralogous organization provides an intrinsic layer of functional redundancy, ensuring robust chromatin remodeling activity even when one component is compromised. Analogous redundancy mechanisms have been observed in other epigenetic regulators, such as the *TET1/2/3* family that regulates DNA methylation^12^. Indeed, only the combined loss of all three *TET* paralogs produces severe developmental phenotypes in mice, while single or double knockouts are largely tolerated^13^. Such compensation is particularly crucial during embryonic development, when precise chromatin regulation is required for correct lineage specification and organism viability. Beyond developmental robustness, paralog redundancy can have therapeutic implications. In cancers driven by mutations in chromatin remodelers, the remaining intact paralog often becomes essential, making it a promising target for selective therapy^14^. Redundancy also exists across the four main chromatin remodeler families which include SWI/SNF, ISWI, CHD, and INO80^15^. Collectively these remodelers maintain chromatin accessibility and transcriptional fidelity^16^. Indeed, yeast studies show that deletion of any single remodeler family produces only modest global effects on chromatin accessibility^17^. Nevertheless, this extensive network of overlapping functions stands in contrast to the strong disease association of specific SWI/SNF paralog mutations in neurodevelopmental disorders. These findings imply that certain paralogs have evolved specialized functions that are critical for neurodevelopment and cannot be compensated by other paralogs.

SWI/SNF plays essential roles in neurodevelopment, with multiple studies highlighting the contributions of its paralogous subunits, including *SMARCA2/4*^18,19^, *SMARCC1/2*^20^, *ARID1A/B*^21,22^, *ACTL6A/B*^23,24^. Although mutations in *SMARCD1*, *SMARCD2*, and *SMARCD3* have been identified in patients with neurodevelopmental disorders^25,26^, the specific functions of SMARCD paralogs during mammalian brain development remain poorly understood. In patients, *SMARCD* mutations are predominately heterozygous missense mutations (84.3%), although nonsense (6.7%), non-frameshift insertions/ deletions (5.6%) and frameshift (3.4%) mutations have also been detected^26^. SMARCD subunits associate with SMARCC proteins to form the initial core of the SWI/SNF complex, serving as essential components required for complex assembly and activity despite lacking ATPase function. In *Drosophila*, a single SMARCD ortholog, *BAP60*, regulates neuronal activity-dependent gene expression programs in the adult brain^25^. In mammals, specific functions for the three SMARCD paralogs have been identified in diverse tissues: SMARCD1 regulates liver metabolism^27^, SMARCD2 controls granulocyte gene expression programs^28^, and SMARCD3 acts as a master regulator of muscle cell fate^29^. All three SMARCD paralogs are expressed in the developing brain, with *SMARCD1* and *SMARCD3* showing much higher expression levels than *SMARCD2* in neural cell types of the developing cortex.

To investigate the redundant and specific functions of SWI/SNF paralogs in the brain, we focused on SMARCD, a core subunit that is encoded by three paralogs. In vivo knockout of either *SMARCD1* or *SMARCD3* individually did not affect mouse cortical development, and combined deletion of *SMARCD1/3* resulted in only modest cortical phenotypes. Using an in vitro model of neuronal differentiation, we systematically deleted all combinations of *Smarcd1/2/3* and found that loss of one or more paralogs was compensated by increased abundance of the remaining paralogs. Remarkably, this compensation occurred post-transcriptionally, through protein stabilization rather than elevated mRNA expression. Through acute degradation approaches, we defined common and paralog-specific target genes in embryonic stem cells and neurons. Notably, SMARCD3 selectively regulated metabolic gene programs that control oxidative phosphorylation in neurons. Furthermore, proteomics analyses revealed transcriptional regulators that interact preferentially with SMARCD1- or SMARCD3-containing complexes, providing a likely mechanism for paralog-specific SWI/SNF targeting. Together, these findings demonstrate that while SMARCD paralogs exhibit substantial redundancy during cortical development, they retain distinct molecular functions in neurons, highlighting how mutations in SMARCD paralogs can lead to neurodevelopmental disorders.

## Results

### Conditional knockout of Smarcd1 or Smarcd3 does not impact mouse cortical development

SMARCD is a core subunit of the SWI/SNF chromatin remodeling complex, required for complex assembly and remodeling activity despite lacking direct nucleosome engagement or ATPase function^30^ (Fig. 1a). Mutations in *SMARCD1/2/3* paralogs have been reported in patients with neurodevelopmental disorders (Fig. 1b), yet their roles in neurodevelopment remain poorly understood. To investigate SMARCD function in the developing cortex, a brain region critical for cognition and frequently perturbed in NDDs, we generated conditional deletions of *Smarcd1* and *Smarcd3* in mice. We focused on these paralogs because they are highly expressed across neural cell types in both human and mouse cortical development, whereas *Smarcd2* shows minimal expression (Fig. 1b). *Smarcd1^fl/fl^* and *Smarcd3^fl/fl^* mice^27,31^ were crossed with *Emx1Cre*^32^ drivers to restrict gene deletion to the embryonic forebrain from E10.5 onward, and knockout efficiency was confirmed by immunostaining (Fig. 1c; Supplementary Fig. 1a-d). We analyzed male and female conditional knockout (cKO; *Emx1Cre^+/-^Smarcd^fl/fl^*) and control (*Emx1Cre^-/-^Smarcd^fl/fl^*) embryos at E14.5 and E18.5. Surprisingly, loss of *Smarcd1* or *Smarcd3* alone did not alter cortical development. The abundance and distribution of apical progenitors (PAX6^+^) and cortical neurons (NEUN^+^) were comparable between cKO and control embryos (Fig. 1d,e and Supplementary Fig. 1e). Similarly, laminar organization at E18.5 was unaffected, with unchanged numbers of deep-layer (TBR1^+^, layer VI) and upper-layer (CUX1^+^, layers II-III) neurons. Consistent with these findings, *Smarcd1* and *Smarcd3* cKO mice were viable, fertile, and phenotypically indistinguishable from littermate controls. Prior studies have found that deletion of other SWI/SNF subunits such as *Smarca4*^18^ or *Actl6a*^23^ severely disrupts cortical neurogenesis and causes perinatal lethality, revealing an essential role for SWI/SNF in neurodevelopment. Therefore, our data suggest that the remaining SMARCD paralogs can compensate for loss of either *Smarcd1* or *Smarcd3* in vivo, maintaining SWI/SNF complex integrity during cortical development.

**Figure 1.**
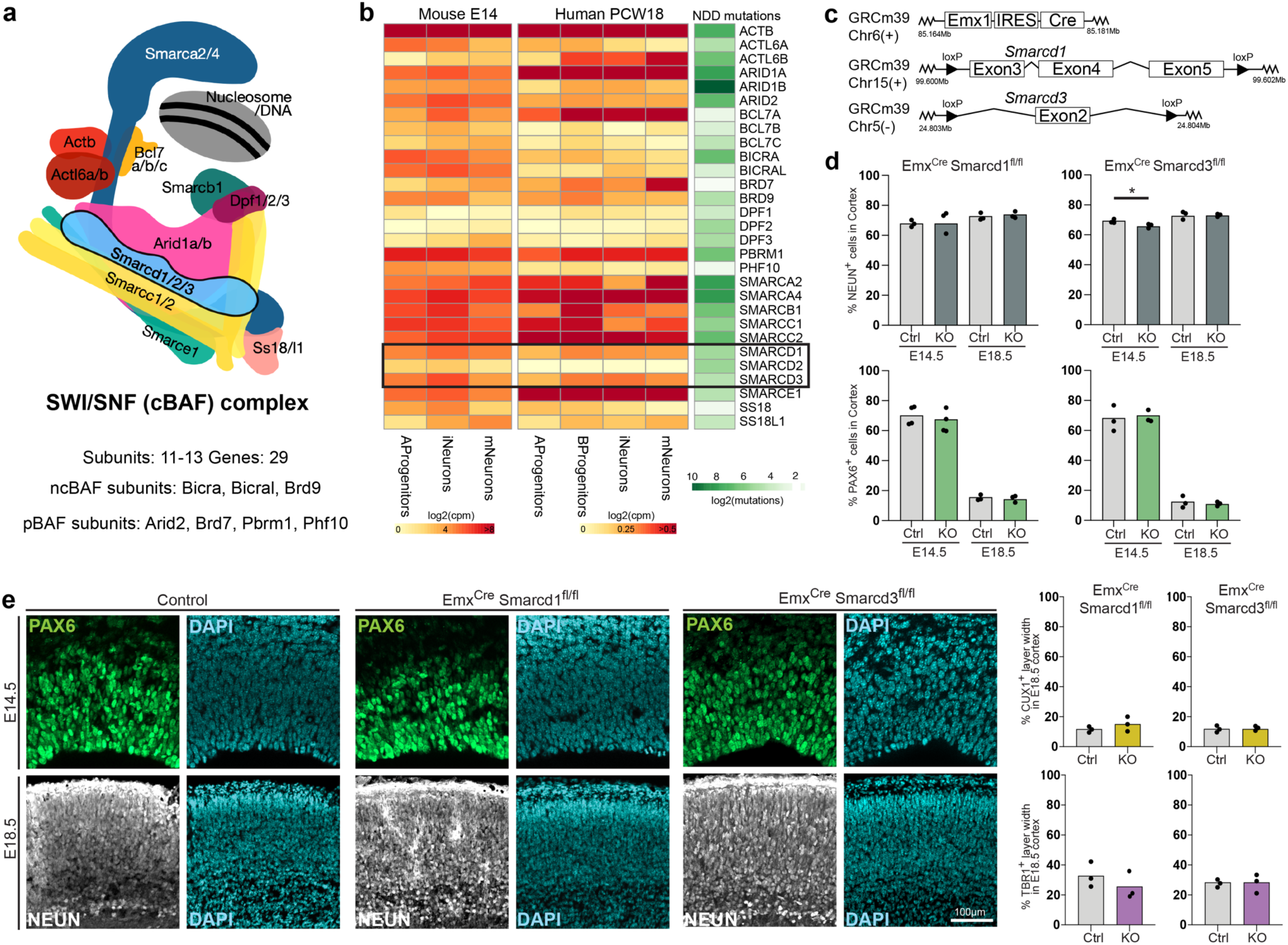
Conditional knockout of *Smarcd1* or *Smarcd3* does not impact mouse cortical development. **a.** Schematic representation of the mammalian SWI/SNF chromatin remodeling complex. **b.** Heatmap showing relative expression levels of SWI/SNF subunits using published scRNA-seq datasets^63^ from mouse E14 and human PCW18 apical progenitors, basal progenitors, immature neurons, and mature neurons, together with their reported mutations in patients with neurodevelopmental disorders^26^ (green). **c.** Schematic representation of the Emx1Cre driver and conditional alleles used for forebrain specific deletion of *Smarcd1* and *Smarcd3*. The Emx1 promoter drives Cre recombinase expression through an internal ribosomal entry site (IRES Cre) cassette, enabling recombination in the developing forebrain. For the conditional floxed *Smarcd1* allele, LoxP sites flank exons 3-5. For the conditional floxed *Smarcd3* allele, LoxP sites flank exon 2. **d.** Quantification of NEUN and PAX6 positive cells in cortical pate and VZ/SVZ respectively, as well as CUX1 and TBR1 positive cell layer width in cortex at E14.5 and E18.5 in Smarcd1 or Smarcd3 cKO embryos. n = 3 embryos per genotype. Data are shown as mean ± SEM. Statistical significance was assessed using a two tailed unpaired Mann Whitney test. **e.** Representative images of coronal brain sections from control and conditional knockout embryos at E14.5 and E18.5. Sections were stained for NEUN and PAX6 at E14.5, and for NEUN, PAX6, CUX1, and TBR1 at E18.5. DAPI was used to visualize nuclei. Scale bars: 100 µm.

### Conditional Smarcd1/3 double knockout mice display mild cortical defects

We next examined the combined requirement for SMARCD1 and SMARCD3 during cortical development. *Emx1Cre^+/-^Smarcd1^fl/fl^* and *Emx1Cre^+/-^Smarcd3^fl/fl^* mice were crossed to generate conditional double knockout (double cKO; *Emx1Cre^+/-^Smarcd1^fl/fl^Smarcd3^fl/fl^*) and control (*Emx1Cre^-/-^Smarcd1^fl/fl^Smarcd3^fl/+^*) embryos. SMARCD1 and SMARCD3 loss was confirmed to be restricted to the forebrain (Supplementary Fig. 2a). Knockout of both *Smarcd1* and *Smarcd3* in the developing cortex produced only subtle cellular phenotypes. The number of apical progenitors (PAX6^+^) was unchanged at E14.5 and E18.5, and only a modest increase in cortical plate neurons (NEUN^+^) was detected at E18.5 (Fig. 2a,b). Laminar identity was preserved, with normal distribution of deep-layer (TBR1^+^) and upper-layer (CUX1^+^) neurons (Fig. 2b, Supplementary Fig. 2b). Despite minimal changes in neuronal composition and layering, cortical thickness was significantly reduced in double cKO embryos at both stages (Fig. 2c,d). Compared to controls, cortical thickness decreased by 28.2% at E14.5 and 33.7% at E18.5, exceeding the small reduction observed in *Smarcd1* single knockouts (15.8%). Nonetheless, double mutants were viable and fertile. These findings indicate that SMARCD1 and SMARCD3 are jointly required for normal cortical growth but are dispensable for neuronal fate specification or laminar organization. This phenotype contrasts sharply with the severe thinning of the cortex, major laminar disorganization and perinatal lethality observed in *Smarca4* or *Actl6a* mutants^18,23^. Thus, our results suggest a partial compensation by SMARCD2 in the developing cortex of *Smarcd1/3* double cKO mice, as at least one SMARCD paralog is required for SWI/SNF complexes to function.

**Figure 2.**
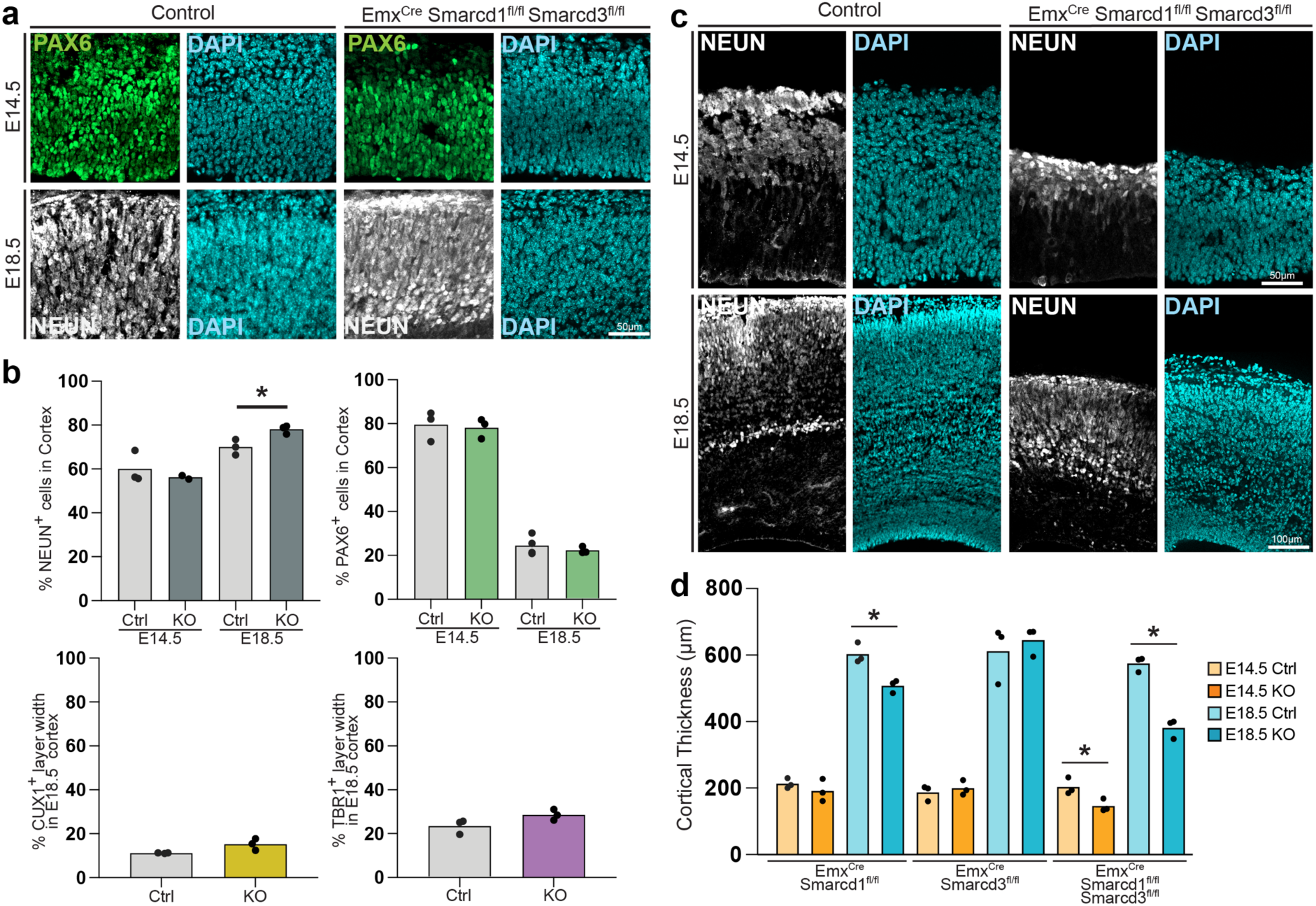
Conditional *Smarcd1/3* double knockout mice display mild cortical defects. **a.** Representative images of coronal brain sections from control and *Smarcd1/3* double knockout embryos at E14.5 and E18.5. Sections were stained for PAX6 at E14.5 and NEUN at E18.5. DAPI was used to visualize nuclei. Scale bars: 50 µm. **b.** Quantification of NEUN and PAX6 positive cells in cortical pate and VZ/SVZ respectively, as well as CUX1 and TBR1 positive cell layer width in cortex at E14.5 and E18.5 in *Smarcd1/3* double knockout embryos. n = 3 embryos per genotype. Data are shown as mean ± SEM. Statistical significance was assessed using a two tailed unpaired Mann Whitney test. **c.** Representative images illustrating cortical thickness differences between Control and *Smarcd1/3* double knockout embryos at E14.5 and E18.5. Sections were stained for NEUN and DAPI. Scale bars: 50 µm (top) and 100 µm (bottom). **d.** Quantification of cortical thickness at E14.5 and E18.5 in *Smarcd1/3* double conditional knockout embryos. n = 3 embryos per genotype. Data are shown as mean ± SEM. Statistical significance was assessed using a two tailed unpaired Mann Whitney test.

### Loss of SMARCDs is compensated by increased stability of remaining paralogs in ESCs

To dissect the molecular basis for SMARCD paralog compensation, we established an in vitro system enabling the generation of all single, double, and triple *Smarcd* knockouts. Using CRISPR/Cas9^33^, we created mouse embryonic stem cell (ESC) lines carrying defined deletions of *Smarcd1/2/3*. For each gene, two sgRNAs targeting exon 4 were used to introduce indels (Fig. 3a). Homozygous editing was confirmed by PCR and Sanger sequencing for all single and double mutants (Supplementary Fig. 3a). Complete loss of all three *Smarcd* genes proved non-viable, phenocopying the deletion of the ATPase subunit *Smarca4* in ESCs^34,35^ and confirming the essential requirement of at least one SMARCD subunit for SWI/SNF complex assembly. Growth and viability assays revealed normal proliferation across most knockout lines (Fig. 3b, Supplementary Fig. 3b), with the exception of *Smarcd1/2* double knockouts, which displayed reduced proliferation, lower viability, and morphological features consistent with premature differentiation (Supplementary Fig. 3c). RNA-seq analysis of *Smarcd1/2* double KO cells confirmed efficient depletion of *Smarcd1* and *Smarcd2* transcripts, while *Smarcd3* mRNA remained unchanged (Fig. 3c). Furthermore, we observed downregulation of pluripotency regulators (*Sox2*, *Esrrb, Dppa5a, Prdm14*) and upregulation of mesodermal genes (*Nodal*, *Bmp4*, *Wnt5a*, *Hox*), consistent with GO-term enrichment for mesoderm differentiation (Fig. 3c,d; Supplementary Fig. 3d-f). These findings align with prior evidence that SMARCD3 promotes cardiomyocyte identity^36^, suggesting that SMARCD3 expressions drives *Smarcd1/2* KO cells toward a mesodermal fate. Interestingly, western blot analysis revealed strong compensatory upregulation of remaining SMARCD proteins across mutant lines (Fig. 3e,f). Loss of SMARCD1 elevated SMARCD2 and SMARCD3 levels, while combined deletion of SMARCD1 and SMARCD3 increased SMARCD2 abundance. In contrast, loss of SMARCD2 or SMARCD3 had little effect on SMARCD1, as it is already the most abundant paralog in ESCs. These data indicate that cells can adjust SMARCD protein levels across paralogs to maintain sufficient SWI/SNF complex assembly. Remarkably, RT-qPCR analysis showed no corresponding changes at the transcript level, as deletion of one paralog did not induce transcriptional upregulation of the others (Fig. 3g). Thus, SMARCD compensation likely occurs post-transcriptionally. Given that unincorporated SWI/SNF subunits are targeted for degradation by E3 ubiquitin ligases^37,38^, our data support a model in which SMARCD paralogs compete for complex incorporation, with incorporation conferring protein stability. When one or more paralogs are lost, the remaining SMARCD subunits gain access to complex assembly and are stabilized, increasing protein levels without the need for increased gene expression.

**Figure 3.**
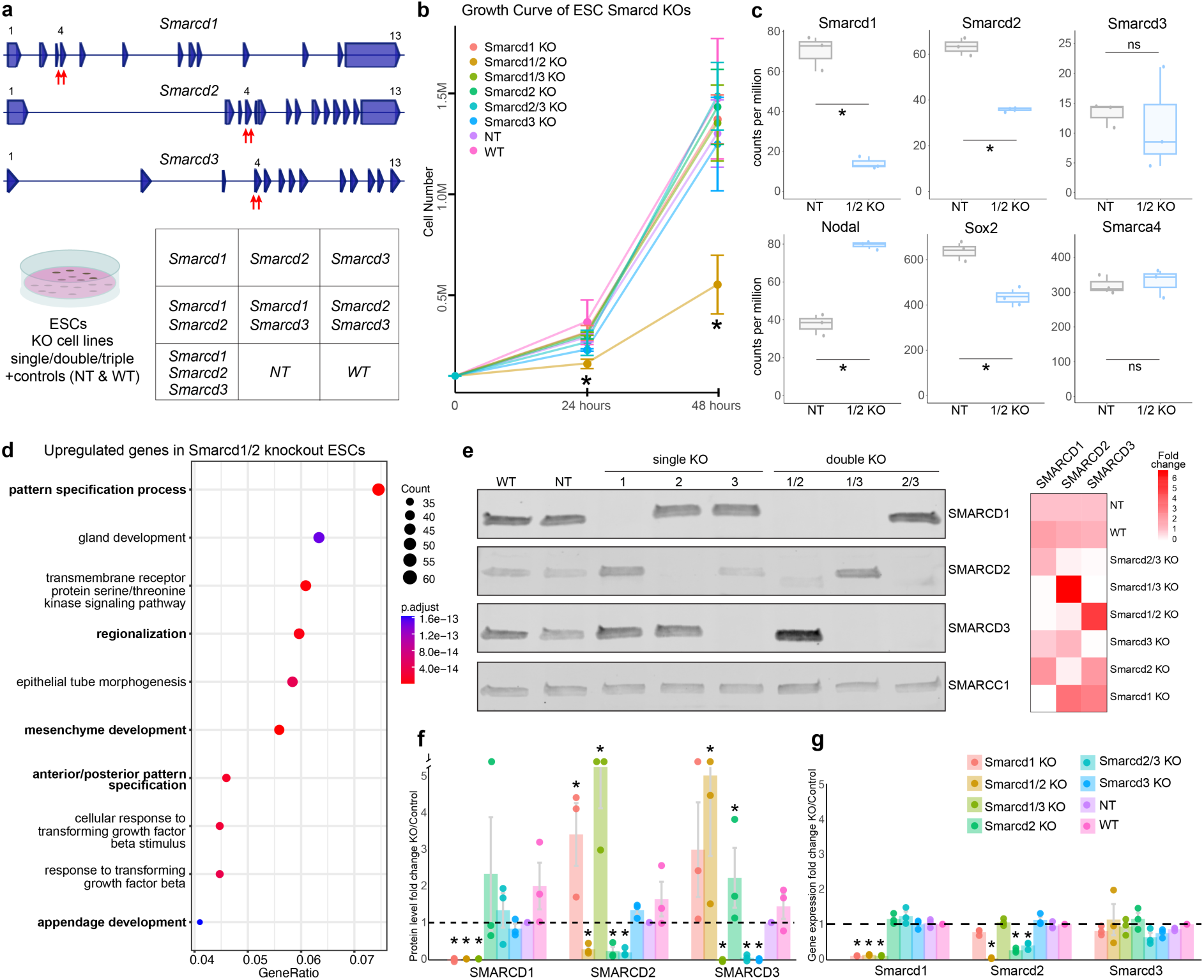
Loss of SMARCDs is compensated by increased stability of remaining paralogs in ESCs. **a.** Schematic of mouse *Smarcd1/2/3* genomic loci with sgRNAs used to target exon 4 for deletion via CRISPR/Cas9 genome editing. Table summarizing the combinations of single, double, and triple KO ESC lines generated. **b.** Quantification of cell numbers (in millions) over 48 hours in culture for all KO cell lines compared to WT and non-targeting sgRNA (NT) controls. Plot reveals normal growth at 24 and 48 hours for all lines, except the *Smarcd1/2* double knockout. n=3. p-value < 0.05 using a two-sided t-test. **c.** RNA-seq analysis of the pluripotency marker Sox2, the mesodermal marker Nodal, the SWI/SNF ATPase subunit Smarca4, and the three Smarcd paralogs in *Smarcd1/2* KO cells compared to NT controls. n=3. mean ± SEM. adjusted p-value < 0.05. **d.** GO term analysis of upregulated genes in *Smarcd1/2* double knockout ESCs. Point size reflects gene count and color indicates adjusted p-values. The x-axis shows the gene ratio. **e.** Western blot showing protein expression of SMARCD1/2/3 in SMARCD single and double knockout ESCs and control cells. SMARCC1 was included as control to assess levels of other SWI/SNF subunits. Heatmap displays fold change in SMARCD levels for each cell line relative to NT control cells. **f.** Quantification of western blots indicates fold change in SMARCD protein levels in single and double knockout ESCs, showing strong compensation of subunit protein levels. n=3. mean ± SEM. p-value < 0.05. **g.** Gene expression analysis of SMARCD subunits by RT-qPCR. Plot shows fold changes in *Smarcd1/2/3* transcript levels in single and double knockout ESCs relative to controls, showing no compensatory changes at the transcript level. n=3. mean ± SEM. p-value < 0.05.

### SMARCD paralog compensation is maintained during neuronal differentiation

To determine whether SMARCD paralog compensation persists during neuronal differentiation, we employed an inducible NGN2 overexpression system to convert ESCs into induced neurons (iNeurons)^39^. Using lentiviral vectors, we stably expressed NGN2-ERT2, a tamoxifen (TAM)-inducible fusion protein that enables temporal control of NGN2 transcriptional activation^40^. In the absence of TAM, NGN2-ERT2 remains cytoplasmic, but following TAM treatment it rapidly translocates to the nucleus to initiate the neuronal differentiation program (Fig. 4a). Seven days after TAM induction, immunostaining confirmed efficient neuronal conversion, with cells expressing TUJ1 and exhibiting characteristic neuronal processes (Fig. 4c, Supplementary Fig. 4f). RNA-seq further validated their neuronal identity: *Rbfox3* and *Dcx* were robustly expressed, while pluripotency markers (*Oct4*, *Nanog*) and glial markers (*Gfap*, *Olig1*) were absent (Fig. 4b; Supplementary Fig. 4b-d). Western blot analysis revealed dynamic changes in SMARCD paralog levels during differentiation (Supplementary Fig. 4a). Furthermore, RNA-seq analysis showed that *Smarcd*1 and *Smarcd*3 expression increased, while *Smarcd2* decreased upon neuronal conversion (Fig. 4d). This expression pattern mirrored published scRNA-seq data from mouse and human cortical development, in which *Smarcd1* and *Smarcd3* are enriched in neurons, whereas *Smarcd2* levels are very low (Fig. 1b). We next generated iNeurons from the SMARCD single and double knockout ESC lines. All mutant lines differentiated efficiently following TAM induction, except for *Smarcd1/2* double knockouts, which failed to acquire neuronal identity, consistent with their prior mesodermal commitment (Supplementary Fig. 4e). Again, western blot analysis confirmed post-transcriptional compensation of SMARCD paralogs in iNeurons (Fig. 4e,f). In *Smarcd1* KO iNeurons, SMARCD2 and SMARCD3 were upregulated; in *Smarcd1/3* double KOs, SMARCD2 levels increased markedly. Interestingly, *Smarcd1/3* double knockouts were competent for neuronal differentiation, suggesting SMARCD2 can compensate for the roles of SMARCD1 and SMARCD3 during neural differentiation, in support of our in vivo findings. In contrast, loss of SMARCD2 had minimal effect on other paralogs, reflecting its low expression in neurons. Together, these findings demonstrate that SMARCD paralogs maintain compensatory regulation during neuronal differentiation, ensuring sufficient incorporation of core SMARCD subunits to sustain SWI/SNF complex assembly and function in postmitotic neurons.

**Figure 4.**
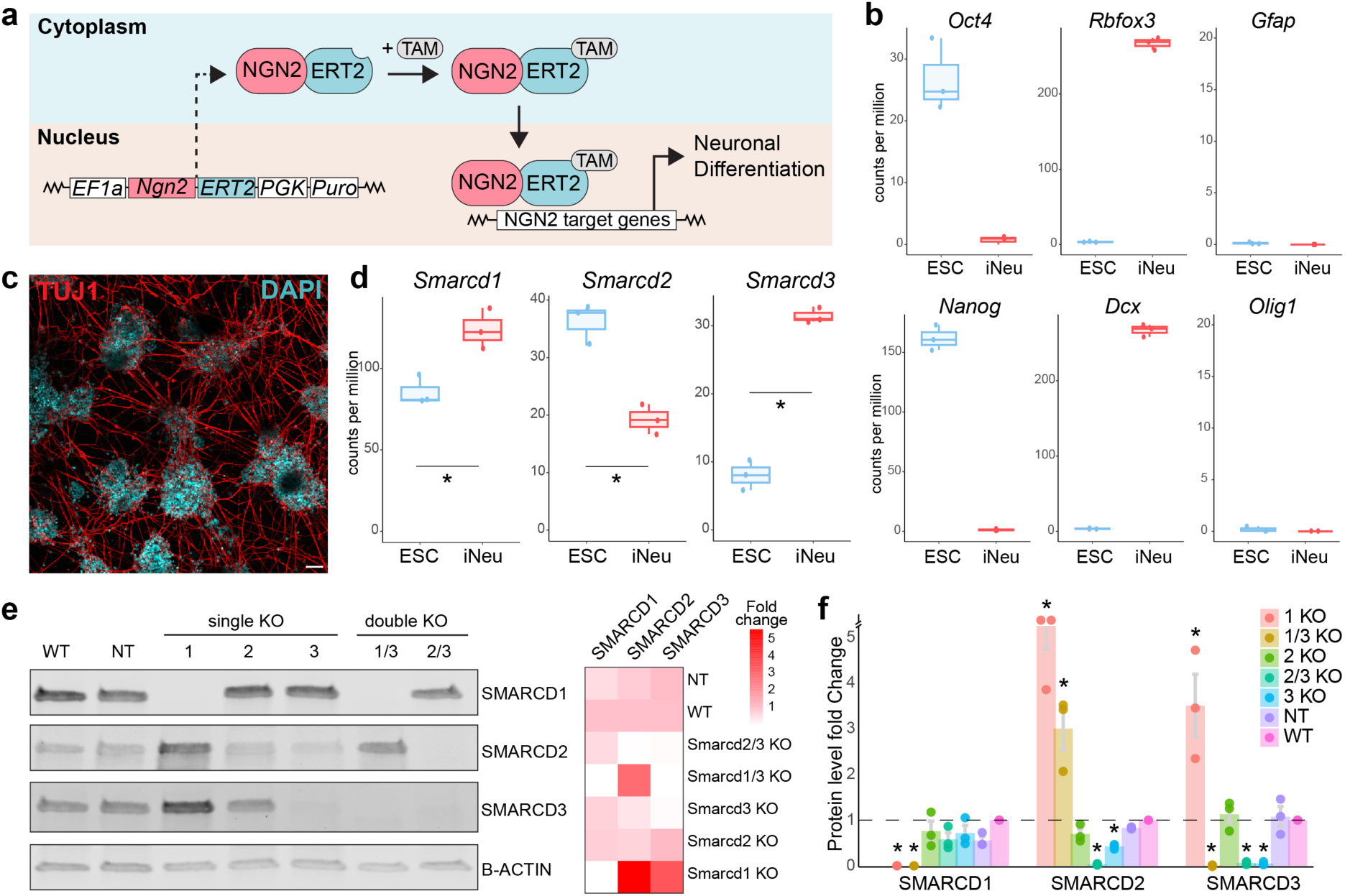
SMARCD paralog compensation is maintained during neuronal differentiation. **a.** TAM-inducible activation of NGN2 drives neuronal differentiation of mouse ESCs. ESCs were transduced with a lentiviral vector expressing NGN2-ERT2. In the absence of TAM (4-hydroxy-tamoxifen), NGN2-ERT2 remains cytoplasmic and inactive. TAM treatment triggers nuclear translocation of NGN2-ERT2 and activation of a pro-neuronal transcriptional program. **b.** RNA-seq analysis of control iNeurons compared to control ESCs. Marker analysis: pluripotency (Oct4, Nanog), neurons (NeuN, Dcx), and glia (GFAP, Olig2). n=3. mean ± SEM. adjusted p-value < 0.05. **c.** Representative image of iNeurons stained for neuronal marker TUJ1 and DAPI. Scale bar: 50 µm. **d.** RNA-seq analysis of control iNeurons compared to control ESCs. All paralogs showed significant differences between ESCs and iNeurons reflecting in vivo kinetics observed in published scRNA-seq data of the developing cortex, see Figure 1b. n=3. mean ± SEM. adjusted p-value < 0.05. **e.** Western blot showing protein expression of SMARCD1/2/3 in *Smarcd* single and double knockout iNeurons and control cells. Beta-actin served as a loading control. Note: a non-specific band was detected in *Smarcd2* single knockout iNeurons. Repeated trials showed the same outcome. The iNeurons were generated from the *Smarcd2* KO ESCs which were validated by genotyping and Sanger sequencing of the mutated *Smarcd2* alleles confirming indels that produce frame-shifts. Heatmap displays fold change in SMARCD levels for each cell line relative to WT control cells. **f.** Quantification of western blots indicates fold change in SMARCD protein levels in single and double knockout iNeurons, showing strong compensation of subunit protein levels. n=3. mean ± SEM. p-value < 0.05.

### Rapid degradation of SMARCDs reveals common and paralog-specific target genes

Although our data indicate robust compensation among SMARCD paralogs, human genetic studies have identified mutations in *SMARCD1/2/3* in patients with neurodevelopmental disorders, suggesting that each paralog may perform distinct, nonredundant functions. To uncover paralog-specific direct target genes and mechanisms beyond compensatory protein stabilization, we employed the dTAG degron system^41^ to achieve rapid and inducible degradation of individual SMARCD proteins (Fig. 5a). Using CRISPR/Cas9, we inserted FKBPv degron and V5 epitope tags at the C-termini of SMARCD1 and SMARCD2, and used a piggyBac strategy to tag the C-terminus of SMARCD3 (Supplementary Fig. 5a-b). The FKBPv-tagged SMARCD protein levels were slightly lower compared to untagged proteins in wild-type ESCs, however the SMARCD1^FKBPv^, SMARCD2^FKBPv^, and SMARCD3^FKBPv^ cell lines displayed normal ESC morphology and growth rates (Supplementary Fig. 5c). Western blot time-course analysis confirmed rapid and efficient degradation upon dTAG-13 treatment. All SMARCD proteins were strongly reduced within 2 h and nearly absent by 6-8 h (Fig. 5b). We next performed RNA-seq in both ESCs and iNeurons at 8 h and 24 h after dTAG-13 treatment to identify direct transcriptional targets of each SMARCD paralog (Supplementary Fig. 5d). Each paralog regulated both shared and distinct sets of genes, supporting common and paralog-specific transcriptional programs (Fig. 5c). Degradation of SMARCD proteins caused both gene upregulation and downregulation, consistent with previous degron-based studies of SWI/SNF subunits (Fig. 5d, Supplementary Fig. 5e). In ESCs, SMARCD1 and SMARCD2 controlled overlapping and unique gene networks linked to chromatin remodeling (ex: *Ino80d, Chd6*, *Chd2, Nfat5*) histone methylation (ex: *Kmt2a*, *Ash1l*, *Kmt2d*) and DNA methylation (ex: *Tet1*, *Tet2*), whereas SMARCD3 loss produced comparatively modest transcriptional changes. In iNeurons, SMARCD1 and SMARCD3 governed distinct metabolic gene programs, while SMARCD2 depletion had minimal impact (Fig. 5e, Supplementary Fig. 6c). Notably, SMARCD3 degradation specifically upregulated a set of genes involved in oxidative phosphorylation, including the mitochondrial electron-transport chain components *Cox5a, Ndufs2, Idh1, Sdhaf4, Ndufa8, Ndufb11, Cox8a, Uqcrq* and signaling kinases *Akt1, Cdk1*; highlighting a role for SMARCD3 in the regulation of neuronal energy metabolism (Fig. 6a).

**Figure 5.**
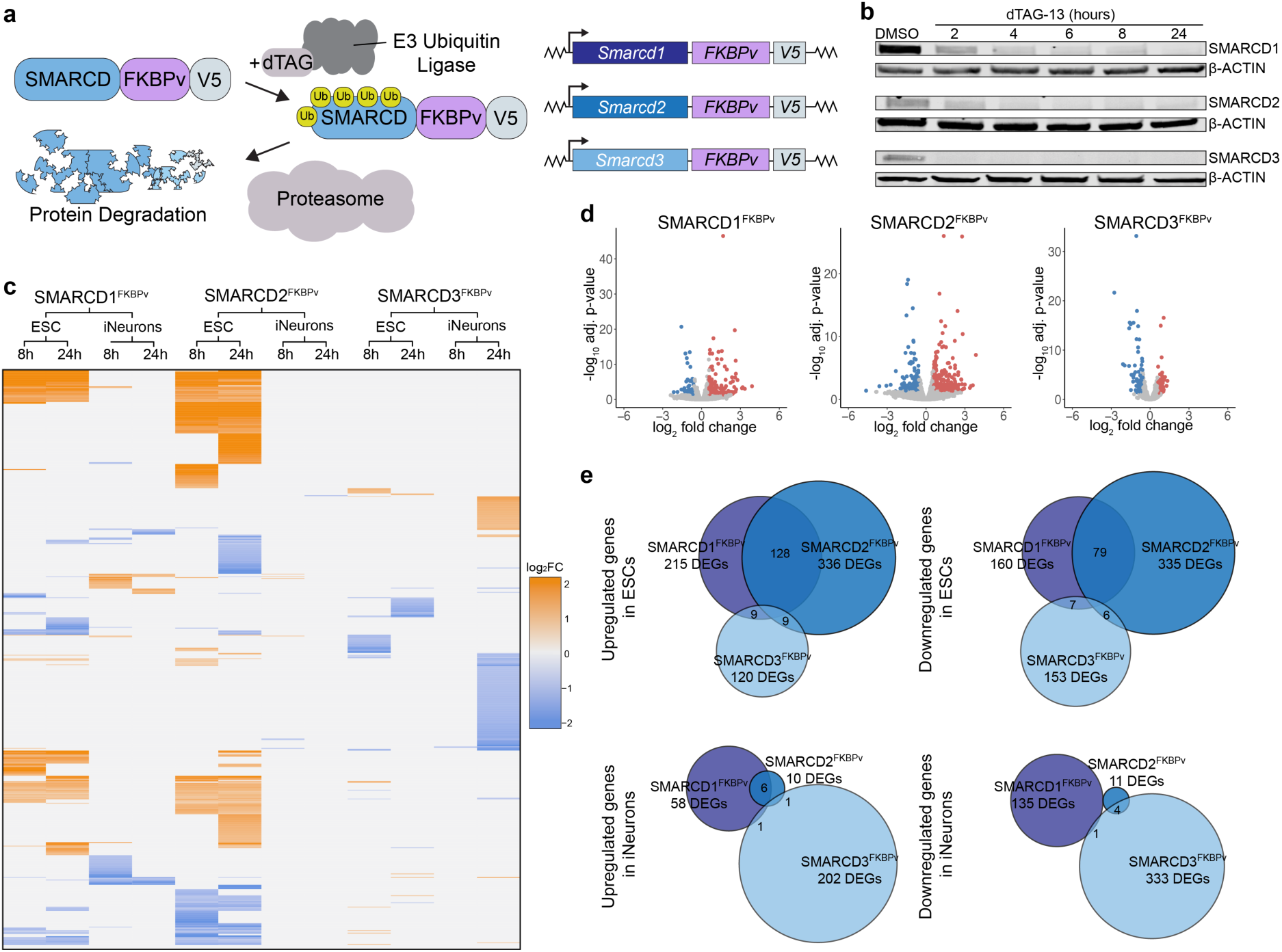
Rapid degradation of SMARCDs reveals common and paralog-specific target genes. **a.** Schematic representation of SMARCD proteins tagged with FKBPv and V5. Trewatment with dTAG-13 recruits an E3 ubiquitin ligase, leading to ubiquitination of SMARCD^FKBPv^ proteins and subsequent proteasomal degradation. **b.** Time course of dTAG-13 induced degradation of SMARCD1, SMARCD2, and SMARCD3 in mouse ESCs. Western blot showing protein levels after treatment with dTAG-13 for 2, 4, 6, 8, and 24 hours. DMSO served as a vehicle control. Beta-actin was used as a loading control. **c.** Heatmap showing differentially expressed genes from RNA-seq analysis of SMARCD^FKBPv^ ESCs and iNeurons after 8 and 24 hours of dTAG-13 treatment. The log fold change scale indicates upregulated genes (orange) and downregulated genes (blue). Each column represents a gene with log two-fold change indicated when statistically significant. n=3. adjusted p-value < 0.05. **d.** Volcano plots depict differentially expressed genes from RNA-seq analysis of SMARCD^FKBPv^ ESC lines after 8 hours of dTAG-13 induced degradation. Each point represents a gene, with log2-transformed fold change on the x-axis and –log10-transformed adjusted P-value on the y-axis. Genes meeting significance thresholds (adjusted P < 0.05 and | FC | > 1.5) are shown in red (upregulated) or blue (downregulated); non-significant genes are shown in grey. **e.** Venn diagrams showing unique and shared upregulated and downregulated DEGs across the three SMARCD^FKBPv^ lines in ESCs and iNeurons at both timepoints.

**Figure 6.**
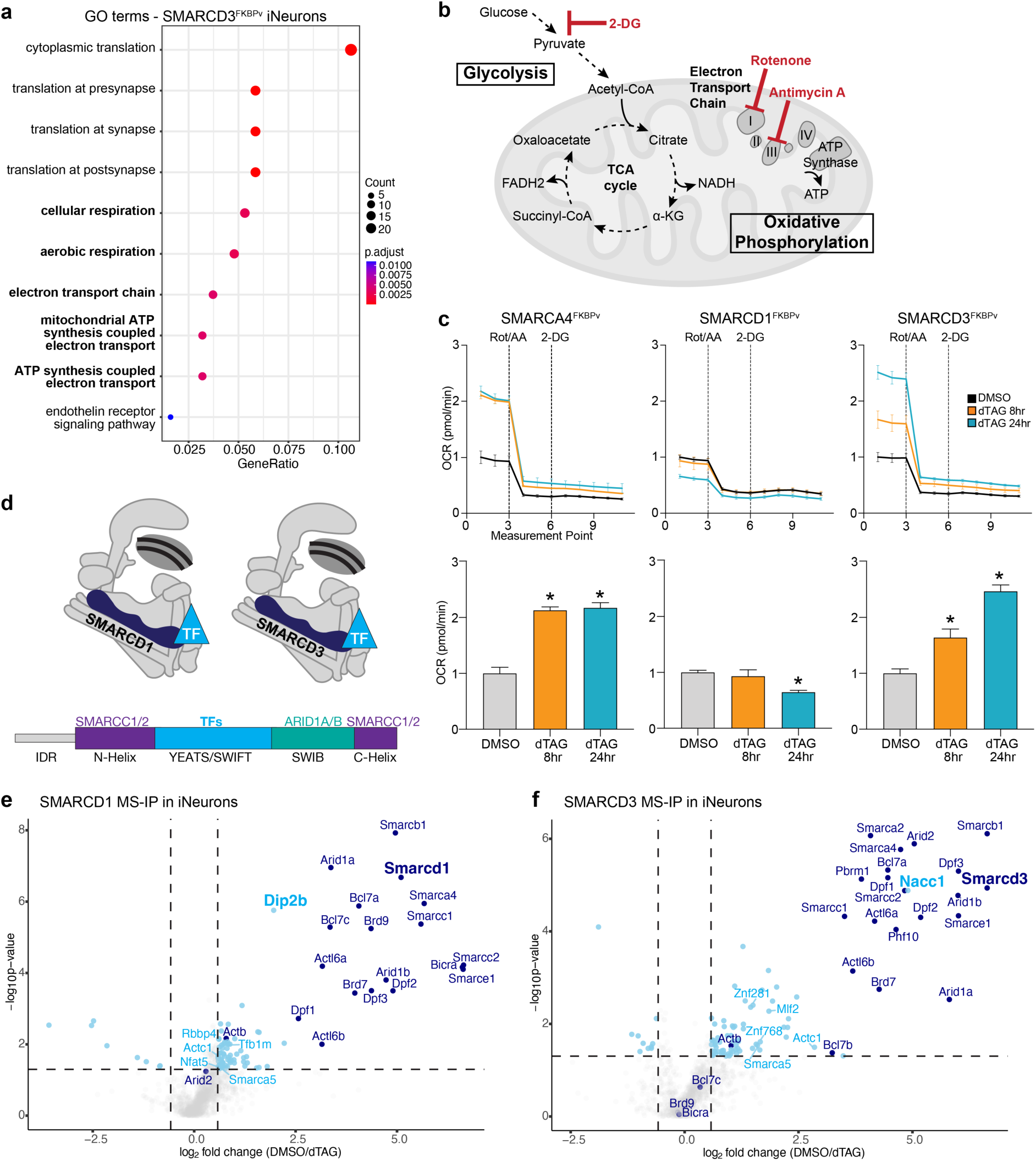
SMARCD3 regulates OXPHOS metabolism in neurons and interacts with a distinct set of proteins. **a.** GO term analysis of differentially regulated genes in SMARCD3^FKBPv^ iNeurons after 24 hours of dTAG-13 treatment reveals a strong enrichment for OXPHOS metabolism genes. **b.** Schematic representation of cellular metabolism, focusing on glycolysis in the cytoplasm and oxidative phosphorylation in mitochondria. In the Seahorse assay, 2-DG blocks glycolysis, while rotenone and antimycin A block oxidative phosphorylation. **c.** Glycolytic rate in SMARCD^FKBPv^ tagged iNeurons following dTAG-13 treatment. The glycolytic rate was measured at 11 sequential time points in SMARCA4^FKBPv^, SMARCD1^FKBPv^, and SMARCD3^FKBPv^ iNeurons. The x-axis shows measurement points 1 to 11, with tick marks at points 3, 6, and 9 (corresponding to 14.2, 33.5, and 52.9 mins, respectively). Cells were treated with dTAG-13 for 8 and 24 hours and compared to DMSO treated controls. Rotenone and antimycin A were injected after measurement 4, followed by 2-DG after measurement 7. Bar plots below show OCR as the average of the first three points prior to rotenone and antimycin A. Data represent mean ± SEM. n ≥ 4. Significant differences relative to DMSO controls are indicated by asterisks (p-value < 0.05). Two-way repeated measures ANOVA was used to evaluate basal effects of dTAG treatment. **d.** Schematic representation of SMARCD subunit domains. The YEATS/SWIFT domain confers specific interactions with TFs. The SWIB domain interacts with ARID1 subunits whereas the N- and C-Helix domains interact with SMARCC subunits. IDRs may also contribute to SMARCD-TF interactions. **e.** Mass spectrometry analysis of SMARCD1^FKBPv^ iNeurons treated with dTAG-13 for 12 hours. The dTAG treated samples were used as the negative controls. V5-trap magnetic particles pulled down all SWI/SNF components (dark blue) and several interactors (light blue). Statistical significance set at p-value < 0.05 and | FC | > 1.5 **f.** Mass spectrometry analysis of SMARCD3^FKBPv^ iNeurons treated with dTAG-13 for 12 hours. The dTAG treated samples were used as the negative controls. V5-trap magnetic particles pulled down all SWI/SNF components (dark blue) and several interactors (light blue). Statistical significance set at p-value < 0.05 and | FC | > 1.5.

### SMARCD3 is required for SWI/SNF-dependent regulation of OXPHOS metabolism in neurons

To functionally assess the metabolic pathways regulated by SMARCD1 and SMARCD3 in neurons and embryonic stem cells, we analyzed cellular metabolism following rapid SMARCD degradation. Stem cells and neurons rely on distinct metabolic programs: neural stem cells primarily use aerobic glycolysis to generate acetyl-CoA for mitochondrial respiration, whereas neurons depend on oxidative phosphorylation (OXPHOS) for ATP synthesis^42^. We measured real-time oxygen consumption rates (OCR; OXPHOS) and extracellular acidification rates (ECAR; glycolysis) using the Seahorse metabolic flux analysis (Fig. 6b). SMARCA4^FKBPv^ cells served as a positive control, as degradation of the SWI/SNF ATPase subunit SMARCA4 broadly disrupts complex activity and metabolic regulation. Indeed, dTAG-13-induced degradation of SMARCA4 produced rapid and pronounced metabolic shifts: in ESCs, both OCR and ECAR decreased by ∼40%, while in iNeurons, ECAR decreased by 38% and OCR increased by 113% (Fig. 6c; Supplementary Fig. 6a,b). These results align with previous studies implicating SWI/SNF in metabolic gene regulation^43^. Strikingly, SMARCD loss phenocopied these changes in a paralog-specific manner. In neurons, OXPHOS activity was selectively dependent on SMARCD3-containing SWI/SNF complexes (Fig. 6c). SMARCD1 degradation had minimal impact, whereas SMARCD3 loss markedly increased oxidative metabolism rates, as observed following SMARCA4 depletion. Together, these results demonstrate that SMARCD3 uniquely regulates the expression and function of OXPHOS-related genes, identifying it as a key SMARCD paralog controlling neuronal OXPHOS metabolism.

### SMARCD1 and SMARCD3 interact with distinct transcription factor networks in neurons

SWI/SNF chromatin remodeling complexes lack intrinsic DNA sequence specificity and instead rely on other chromatin regulators and transcription factors (TFs) for genomic targeting^44,45^. Prior studies have shown that SMARCD subunits mediate such interactions in various cell types, and structural analyses identified the YEATS/SWIFT domain of SMARCD proteins as a key platform for SWI/SNF-TF binding^46^ (Fig. 6d). To identify paralog-specific partners in neurons, we performed immunoprecipitation-mass spectrometry (IP-MS) of SWI/SNF complexes containing either SMARCD1 or SMARCD3, using V5-Trap nanobody pulldowns in control and dTAG-13 treated iNeurons. Pulldowns efficiently recovered all 11 canonical SWI/SNF subunits, validating our IP-MS approach. Beyond dedicated complex subunits, we identified 73 interactors of SMARCD1-containing complexes and 75 interactors of SMARCD3-containing complexes. Most were unique transcriptional regulators (ex: NFAT5, RBBP4, ZNF281, ZNF768), with a subset common to both complexes (SMARCA5, DDX39A, and ACTC1). Notably, the most enriched interactors, comparable in abundance to core SWI/SNF components, were DIP2B for SMARCD1 complexes and NACC1 for SMARCD3 complexes (Fig. 6e,f; Supplementary Fig. 7a,b). Both DIP2B and NACC1 are known regulators of neuronal gene expression, previously shown to interact with TFs as well as DNA methyltransferases and histone deacetylases, respectively^47–51^. Importantly, mutations in *DIP2B* and *NACC1* also cause neurodevelopmental disorders resembling those linked to *SMARCD* mutations, suggesting shared regulatory pathways^47,52^. Together, these findings reveal that SMARCD1- and SMARCD3-containing SWI/SNF complexes engage distinct transcriptional regulator networks in neurons: in particular SMARCD1 likely controls gene regulation through DIP2B, while SMARCD3 specificity is mediated via its interaction with NACC1.

## Discussion

Although chromatin regulators operate in all cell types, mutations in these pathways are disproportionately associated with neurological phenotypes. More than 40 mendelian disorders are linked to chromatin regulators, and over 90% present with neurodevelopmental features^6^. Thus, while mutations in these genes can produce multi-tissue symptoms, their most consistent impact is on the brain. To explore this brain-specific vulnerability, we focused on SMARCD, a core subunit of the SWI/SNF chromatin remodeling complex. Three SMARCD paralogs are expressed in the brain, and all have been implicated in NDDs^26^. Notably, *SMARCD1* de novo mutations cause a syndromic NDD with intellectual disability and distal limb anomalies, phenotypically resembling Coffin-Siris syndrome, which also results from mutations in SWI/SNF subunits^25^.

In contrast to several other SWI/SNF subunits with multiple paralogs, our data reveal that loss of an individual SMARCD paralog can be compensated for during mouse cortical development. We find that deletion of paralogs in embryonic stem cells or neurons increases protein levels of the remaining paralogs without altering their mRNA expression. Similarly, protein level compensation of SMARCD paralogs has also been observed in SWI/SNF mutant HAP1 human cells^53^. This compensation differs from transcriptional adaptation mechanisms described in other contexts^54^, such as *DMD*/*UTRN* paralog compensation in Duchenne muscular dystrophy, which relies on mRNA decay-dependent transcriptional upregulation^55^. We propose that SMARCD compensation occurs at the level of SWI/SNF complex assembly. Prior work showed that incorporation of SMARCB1 into the complex via SMARCC1 prevents its ubiquitination by blocking binding of SMARCB1 to TRIP12, an E3 ubiquitin ligase^38^. Analogously, we suggest that SMARCD paralogs compete for complex incorporation, where incorporation confers protein stability. Loss of one paralog frees assembly sites, allowing stabilization of the remaining subunits without transcriptional changes. Future experiments targeting the proteasome or specific ubiquitin ligases will clarify the mechanisms governing SMARCD stoichiometry and turnover. Paralog redundancy, however, is context-dependent. In hematopoiesis, *Smarcd2* deletion disrupts granulocytic differentiation and cannot be rescued by *Smarcd1* overexpression^28^. Consistent with this, *SMARCD2* loss-of-function mutations underlie neutrophil-specific granule deficiency, an autosomal recessive blood disorder^28^. Together, these findings highlight how tissue-specific requirements for SWI/SNF paralogs shape the diverse phenotypic outcomes of chromatin remodeling defects in human disease^56^.

Using the dTAG degron system, we defined shared and paralog-specific transcriptional programs regulated by SMARCD subunits in mouse embryonic stem cells and neurons. Notably, SMARCD3 controlled a network of genes involved in oxidative phosphorylation in neurons. Importantly, we show that loss of SMARCD3 in neurons has functional consequences for cellular metabolism, as evidenced by increased oxygen consumption rates. This finding aligns with earlier reports showing that SMARCD1 and SMARCD3 modulate metabolic pathways in liver and pancreas, respectively^57–59^. Supporting these observations across species, genetic screens in *C. elegans* identified the *SMARCD* homolog *ham-3* as essential for serotonin metabolism and serotonergic neuron specification^60^. Together, these results underscore a conserved role for SMARCD proteins in neuronal metabolic gene regulation. Furthermore, previous work revealed that the SWI/SNF subunit BCL7A regulates mitochondrial metabolism, a function required for proper neural progenitor fate decisions in mice^43^. Because neurons depend heavily on oxidative phosphorylation for energy^61^, future in vivo analyses of *Smarcd* conditional knockouts may yet reveal functional deficits in neuronal excitability or synaptic activity. These mechanistic links could explain the cognitive impairments observed in individuals with *SMARCD* mutations.

Targeting of SWI/SNF complexes to specific genomic loci depends on interactions with transcription factors, other chromatin regulators, and histone modifications. Several studies have demonstrated direct binding of SMARCD subunits to TFs such as MYOD^29^, C/EBPε^28^, p53^62^, USF1^58^, FOXA1^59^ and PU.1^46^. Recent work identified the SWIFT domain within SMARCD subunits as a versatile binding platform for transcriptional regulators within the SWI/SNF complex. In cancer cells, a single amino acid substitution in this domain disrupts the PU.1-SWI/SNF interaction, impairing site-specific chromatin remodeling^46^. Our data reveal that *SMARCD* paralogs engage distinct protein networks in neurons, thereby potentially conferring specificity to SWI/SNF targeting depending on the incorporated paralog.

For example, we identify DIP2B as a SMARCD1 interaction partner and NACC1 as a SMARCD3 interaction partner. Intriguingly, mutations in *DIP2B* and *NACC1* cause neurodevelopmental syndromes that phenocopy *SMARCD*-related disorders, suggesting convergence on shared regulatory pathways^47,52^. In both neurodevelopmental and oncogenic contexts, a central therapeutic challenge is to achieve cell type-specific modulation of chromatin remodeling. The unique binding interfaces defined by each SMARCD paralog, which may extend beyond the SWIFT domain to IDRs as well, offer a potential route to selective intervention. By targeting the interaction surfaces that mediate paralog-specific SWI/SNF recruitment, it may be possible to fine-tune transcriptional activity in disease-relevant cell types without globally perturbing chromatin architecture.

In summary, SMARCD paralogs act largely redundantly during cortical development, yet our degron and proteomic analyses reveal both common and paralog-specific functions in neurons. These findings expose a dual logic of redundancy and specialization that governs SWI/SNF activity and provide a mechanistic framework for the selective vulnerability of *SMARCD* genes in neurodevelopmental disorders. Furthermore, this study offers insight into why mutations in SWI/SNF components disproportionately affect the brain. In certain tissues, the built-in redundancy among paralogs may be sufficient to buffer decreased levels of a single paralog. However, the prolonged timeline of neuronal maturation and the extraordinary cellular diversity required for brain development may push this compensatory system to its limits, revealing paralog-specific functions that, when disrupted, manifest as neurodevelopmental disease.

## Materials and Methods

### Animal experiments

All mouse experiments for this study were conducted at the University of Geneva, in accordance with the Cantonal Veterinary Office of Geneva (licenses GE188 and GE515). B6.Cg-Smarcd1^tm1.1Jddl^ Emx1^tm1(cre)^ and B6.Cg-Smarcd3^tm1.1Jddl^ Emx1^tm1(cre)^ mice were bred after being obtained from the Jackson Laboratory. Genotypes were determined with the primers listed in Supplementary Table 1.

### Mouse Embryonic Stem Cells

ESCs derived from blastocysts of mixed 129-C57Bl/6 background were cultured in media containing DMEM (Gibco, 31966-021), supplemented with LIF (in-house production), 15% FBS (Gibco, 10437-028), 0.1 mM β-Mercapthoethanol (Gibco, 31350010), 1x non-essential amino acids (Gibco, 11140035), 100 mM HEPES buffer (Gibco, 15630056), 100 U/ml Penicillin-Streptomycin (Gibco, 15140122) at 37°C and 5% CO_2_. All cell culture medium was filtered using the Stericup Filter system (Millipore, S2GPU05RE). Cells were cultured on 10cm plates (Thermoscientific; 172931) coated for 30 min with 0.2% gelatin solution (Sigma, G1890-100G) and passaged every two to three days by dissociation with 0.05 % Trypsin-EDTA (Gibco, 25300054).

### Plasmid cloning

For the dTAG ESC lines, the FKBPv-V5 tag was obtained from Addgene plasmid #91798 and knocked into the *Smarcd1* and *Smarcd2* loci using HDR templates containing 0.5-1 kb homology arms flanking the C-terminal stop codons. The genomic sequences for HDR templates were ordered as gene fragments (Twist Bioscience) using the mm10 genome. The sgRNAs were cloned into Addgene plasmid #62988 that also expresses Cas9. For *Smarcd3*, the cDNA was ordered as a gene fragment (Twist Bioscience) and then fused with an FKBPv-V5 tag and cloned into a piggyBac Addgene plasmid #96930 with a DOX-inducible promoter using In-Fusion HD cloning (Takara, 638947). For dTAG induced protein degradation, cells were treated with 500nM dTAG-13 (Sigma, SML20601-5MG). For the NGN2-ERT2 lentivirus construct, the fusion protein was cloned into Addgene plasmid #102881 using In-Fusion HD cloning of template DNA sequences obtained from Addgene plasmids #52047 and #13777.

### CRISPR/Cas9 genome editing

ESCs were plated on gelatin coated plates with antibiotic resistant DR4 mouse embryonic fibroblasts (MEFs). For CRISPR/Cas9 knock-in a transfection mix containing 4 µg template DNA, 8 µg sgRNA/Cas9 plasmid (Addgene 62988), 200 µl Opti-Mem (Gibco, 11058021) and 25 µl Lipofectamine 2000 (Invitrogen, 11668019) was prepared. For CRISPR/Cas9 knockout a transfection mix containing 4 µg sgRNAa/Cas9 plasmid, 4 µg sgRNAb/Cas9 plasmid (Addgene 62988), 200 µl Opti-Mem and 25 µl Lipofectamine 2000 was prepared. The transfection mixes were incubated for 15 min at RT before adding them to the cells. After 4 h, cell culture media was renewed. After 24 h, cells were put into 1.5 µg/ml Puromycin (InvivoGen, ANT-PR-1) selection for 48 h. Single colonies were manually picked after 3 days, dissociated with trypsin and expanded on MEFs. Feeder free cultured lines were expanded for DNA and protein extraction. ESC lines containing homozygous insertions were confirmed by PCR, Sanger sequencing and western blotting. sgRNA and genotyping primer sequences are provided in Supplementary Table 1.

### piggyBac stable cell lines

ESCs were plated on gelatin coated plates with MEFs and transfected with a mix of 5 µg template plasmid and 2 µg piggyBac transposase plasmid, 200 µl Optimem (Gibco, 11058021) and 25 µl Lipofectamine 2000 (Invitrogen, 11668019). The transfection mix was incubated for 15 min at RT before adding it to the cells. After 4 h, cell culture media was renewed. After 48 h, cells were put into Hygromycin (Invivogen;ant-hg-1) selection for 4 days. Next, single colonies were picked to generate clonal cell lines. Successful transposition and stable expression of DOX-inducible SMARCD3-FKBPv-V5 clones was confirmed via western blot.

### NGN2-derived neuronal differentiation of ESCs

Smarcd^FKBPv^ ESCs were infected with the NGN2-ERT2 expressing lentiviruses and selected for with 1.5 µg/ml Puromycin (Invivogen; ant-pr-1) for 48 hours. For neuronal differentiation, cells were cultured in neurobasal media containing DMEM/F12 (Gibco;31331-028), Neurobasal (Gibco;21104-049), B2 (Gibco; 17504-044), N27 (Gibco;17502_048), NEAA, ß-ME, HEPES and treated with 1µM TAM (4-hydroxy-tamoxifen; CalBioChem;579002) for 48 hours to induce NGN2 activity. The cells were then dissociated with Accutase (Gibco; A11105-01) and plated on Geltrex (Thermofisher; A1413201) coated plates in neurobasal media containing BDNF (Peprotech; 450-02-10UG) and NT-3 (Peprotech; 450-03-10UG). After 7 days of culture, the differentiated iNeurons were analyzed.

### Lentivirus production

For lentivirus production HEK293T cells were seeded on 15 cm cell culture plates in DMEM media (Gibco, 31966-021), supplemented with 10% FBS (Gibco, 10437-028) and 100 U/ml Penicillin-Streptomycin (Gibco, 15140122). Cells were cultured at 5% CO_2_ and 37°C. For transfection, a transfection mix containing 1.8 ml Opti-Mem (Gibco, 11058021), 140 µl PEI (Polysciences, 24765), 4.5 µg plasmid pMD2.G (Addgene 12259), 13.5 µg plasmid psPAX2 (Addgene 12260) and 18 µg of lenti vector DNA was mixed and incubated for 10 min at RT. 1.8 ml of transfection mix was added dropwise to HEK293T cells. The following day the media was changed and two days later, the supernatant was collected and centrifuged at 1000 g for 5 min. The supernatant was filtered and concentrated with Lenti-X (Takara, 631231). The collected virus was resuspended in PBS for infection of ESCs.

### Cell Growth assay

To measure cell growth, 10’000 ESCs were plated in each well of a 6-well plate (thermoscientific; 140675). At two timepoints (24 hours and 48 hours), cells were dissociated using Trypsin-EDTA (0.05%) for 5 minutes at 37 °C. Cells were resuspended in warm cell culture media, and cell number was quantified using a TC20 automatic cell counter (Bio-Rad) after staining with Trypan Blue (Gibco, 15250061).

### ATPlite cell viability assay

The method was performed as previously described by Schwaemmle et al. (2025). 10’000 ESCs were plated in each well of a 96-well plate (Falcon; 353072). After 24 hours, ATPlite reagent (PerkinElmer, 6016943) was added directly to each well containing 50 µL of ESC medium. The plate was shaken for 2 minutes to ensure proper mixing, followed by a brief centrifugation. Luminescence was measured using the SpectraMax L 384-well plate reader (Molecular Devices).

### Mouse brain cryosections

Pregnant mice were anesthetized with isoflurane (Piramal; G22B22A), followed by cervical dislocation and exsanguination. Embryos were dissected at two stages, E14.5 and E18.5, and brains were postfixed with 4% PFA (Biochem; A3813) at 4°C overnight. Brains were stored in 30% sucrose (Milipore; 84100) at 4°C. Brains were frozen in OCT cryoembedding matrix (Milestone; 51420) at -60 °C for 1 minute using the Prestochill cryoembedding system (Milestone) and cut into 30μm thick sections using a cryostat (Leica CM3050 S) at TO 16 °C and TC 20 °C. Brain sections were collected on glass slides (epredia; J1800AMNZ) and left to dry at 37 °C for 10 minutes, and stored at -20 °C.

### Immunohistochemistry

Brain sections were washed in 1x PBS for 5 minutes and incubated for 1 hour in blocking solution (1x PBS; 5% Goat serum; 0.25% Triton X-100). The following primary antibodies were used overnight at 4°C in blocking solution: rabbit anti-NEUN (1:200 dilution; Proteintech); rabbit anti-PAX6 (1:200 dilution; Biolegend); rabbit anti-CUX1(1:200 dilution; Proteintech); mouse anti-TBR1 (1:100 dilution; Proteintech). For CUX1 and TBR1 staining, brain sections were subjected to an antigen retrieval step before the incubation with primary antibodies. Sections were boiled in citrate buffer pH6 (BioChemica; A3901) in a pressure cooker for 20 min and chilled to room temperature for 40 min. After the primary incubation, brain sections were washed in 1x PBS for 10 min, 3 times at RT followed by incubation for 90 min at RT with the following secondary antibodies in blocking solution: anti-rabbit Alexa fluo 488 (1:500 dilution; Lifetech); anti-mouse Alexa fluo 488 (1:500 dilution; Lifetech); anti-rabbit Alexa fluo 568 (1:500 dilution; Lifetech); anti-mouse Alexa fluo 568 (1:500 dilution; Lifetech). Brain sections were washed in 1x PBS for 10 min, for 3 times at RT and nuclear stained with 4′,6-diamidino-2-phenylindole (DAPI; 1:2000 dilution in 1x PBS) for 10 min. Brain sections were washed in 1x PBS for 10 min, for 3 times in RT and dried in dark for 30 minutes. Brain sections were mounted with glass coverslips using Fluoromount-G™ Mounting Medium (SouthernBiotech; 0100-01) and left to dry at 4 °C overnight. Antibody list provided in Supplementary Table 1.

### Imaging and Quantification

For quantification of NEUN, PAX6, CUX1, and TBR1 immunostaining, sections were imaged using an Axiocam Fluorescence 702 mono microscope (Zeiss; Apotome 2; Plan Apochromat 10X/0.45 (WD=2.1mm); EC Plan-Apochromat 20X/ 0.8 (WD=0.55mm); Zen 2.6). For high representative images, a confocal microscope was used: STELLARIS 5 microscope (Illuminator HXP120; two Hybrid S detectors (HyD S1 and HyD S2); HC PL APO CS2 20x/0.75 and HC PL APO 40x/0.95; LAS X software (version: 4.5.0.25531)). For imaging fixed cultured cells, the Zeiss Axio Observer Z1 microscope was used (with Definite Focus 2; EC Plan-Neofluar 10x / 0.30, Ph1 WD 5.2 mm). Quantification of the number of positive cells for each immunostaining was done using Qupath (ver. 0.5.1 on HP Z640) with a custom program script (Nicolas Liaude, Bioimaging facility UNIGE). Data were analysed and plotted using GraphPad Prism 9 (ver. 9.2.0). Illustrative images were edited with ImageJ2 (ver. 2.9.0/1.53t).

### RNA sequencing

For RNA seq, 1 million cells were collected using trypsin, washed in PBS, and stored at -80°C. RNA was extracted from cell pellets using RNeasy kits (Qiagen, 74104). RNA libraries were prepared using Collibri 3’ mRNA Library Prep Kit (Invitrogen, A38110024) or QuantSeq 3’ mRNA Library Prep Kit (Lexogen, 192.96). Library quantification and quality were assessed by Qubit and Tapestation (DNA High sensitivity chip). Libraries were sequenced on an Illumina NovaSeq 6000 sequencer with single-end 100 settings.

### Mass Spectrometry

Cultured neurons were collected using Accutase (Gibco; A11105-01) for 7 min at 37°C. Cells were resuspended in neurobasal media, centrifuged at 500rcf for 5 min. After removing supernatant, cell pellets were resuspended in nuclear extraction buffer (20mM HEPES (Sigma; H3375), 10mM KCl (Sigma;60130), 0.1% Triton X-100 (BioChemica; A1388), 20% Glycerol (Sigma;49767), Protease inhibitor (Roche;11836170001), H2O) and left on ice for 10 min. After incubation, nuclei pellets were resuspended in RIPA buffer (10mM Tris-HCl (Invitrogen; 15504-020) pH8, 150mM NaCl (Sigma;71376), 1mM EDTA (BioChemica; A1104) pH8, 0.5mM EGTA (Sigma; E3889) pH8, 1% NP-40 (BioChemica; A1694), 0.5% DOC (Sigma; D2510), 0.1% SDS (A3942), protease inhibitors, Benzonase (E1024-25KU) and left on ice for 20 min. Pellets were centrifuged at 13,000rcf for 10 min at 4°C. 20µl samples were taken for Western blot as input proteins. After centrifugation, 250µl dilution buffer (1M Tris-HCl pH 7.5, 0.5M EDTA, protease inhibitors) was added to 250µl supernatant. Samples were added to 25µl activated and washed V5 magnetic particles (ChromoTek V5-Trap® Magnetic Particles M-270) and kept on a tube rotating mixer overnight at 4°C. On the following day, samples were placed on a magnet and supernatants were taken for Western Blot as non-bound proteins. After few minutes of magnetic separation, beads were well resuspended in 500µl wash buffer followed by magnetic separation for a total of three washes. After the washes, beads were resuspended in 60µl cold PBS and stored at 4°C. 10µl of bead suspension was taken for Western blot as immunoprecipitated protein. Mass spectrometry analysis was done at the Proteomics core facility of UNIGE. Proteins were on-bead digested with a Trypsin/LysC mix using iST kits (Preomics) and peptides were analysed by nanoLC-ESI-MSMS using a Vanquish NEO liquid chromatography system (Thermo Fisher Scientific) coupled with an Orbitrap Fusion Lumos Mass Spectrometer (Thermo Fisher Scientific). Database searches were performed with Mascot (Matrix Science) using the mouse reference proteome database (Uniprot), including the custom bait SMARCD-FKBPv-V5 protein sequence. Data were analysed and validated with Proteome Discoverer (Thermo Fisher Scientific) with 1% of protein FDR and at least 2 unique peptides per protein. Western blot samples were boiled at 95°C for 5 minutes in 2x SDS-Laemmli sample buffer and DTT and stored at 4°C.

### Seahorse glycolysis and mitochondrial respiration analysis

The day before the metabolic assay of ESCs, 20’000 cells were plated in each well of a Seahorse XF Pro M Cell Culture Microplate (Agilent) and the XF Sensor Cartridge (Seahorse XFe24/XF Flex, Agilent) was hydrated in molecular-grade water overnight at 37°C in a non-CO_2_ chamber. The following day, cells were washed in 1x PBS and 180µl XF assay Medium for ESCs (1x XF assay DMEM medium pH7.4, 15% Fetal Bovine Serum, 1x non-essential amino acids, 1x Penicillin-streptomycin, 100µM ß-mercaptoethanol, LIF (homemade), Glucose (Gibco; A24940), Glutamine (Sigma; 56-85-9), Sodium pyruvate (Gibco;11360-039)) was added to each well. For metabolic assay of cultured iNeurons, XF assay Medium for neurons (1x XF assay DMEM medium pH7.4, 0.5x N-2 supplement (Gibco), 0.5x B-27 supplement (Gibco), 1x non-essential amino acids, 1x Penicillin-streptomycin, 100µM ß-mercaptoethanol, LIF (homemade), Glucose, Glutamine, Sodium pyruvate) was used. Cell culture microplate and XF Sensor Cartridge (molecular grade water was replaced by 200µl of XF Calibrant Solution (Agilent)) were incubated in a non-CO_2_ chamber at 37°C for 1 hour. After incubation, 1.5µM of Rotenin/Antimycin and 50mM of 2-DG (Seahorse XF Glycolytic Rate Assay Kit (Agilent)) were loaded to each well of XF Sensor Cartridge in Port A and Port B respectively. Port C and D were filled with 1x PBS. Cell culture microplate and XF Sensor Cartridge were loaded to Seahorse XFe96 analyser.

### RT-qPCR

RNA was extracted from cell pellets using RNeasy kits (Qiagen, 74104). 500ng RNA was used to make cDNA libraries with PrimeScript RT kits (Takara, RR037A). For the RT-qPCRs, 1 µl of diluted cDNA was mixed with 10 µl of SYBR green (applied Biosystems, A25742), 5 µl of H2O and 4 µl of forward and reverse primers at 2.5 µM. The 96-well qPCR plates were measured on a QuantStudio1 thermocycler (Applied Biosystems). RT-qPCR primer sequences provided in Supplementary Table 1.

### Western blot

Cell pellets were lysed in RIPA lysis buffer (10 mM Tris-HCl pH8, 150 mM NaCl, 1 mM EDTA pH8, 0.5 mM EGTA pH8, 1% NP-40, 0.5% DOC, 0.1% SDS) for 20 min at 4°C. Lysates were centrifuged at 4°C, 13.000 rcf for 10 min and supernatant transferred to new tubes. Protein concentration was determined via BCA protein assay (Pierce, 23225), according to the manufacturer’s instructions. Samples were mixed with 1x Loading buffer (stock at 5x: 10% SDS, 0.2 M Tris HCl pH6.8, 30% Glycerol, Bromophenol blue), 20x DTT (1M) and boiled for 5 min at 95°C. Samples were loaded onto 4-15% Mini-protean TBX gel (BioRad, 4561084) and run for 1 h at 150 V in Running buffer (0.0125 M Tris, 0.096 M Glycine). Transfer was conducted using the Trans-Blot Turbo RTA kit (BioRad, 1704270) according to the manufacturer’s instructions. For total protein staining, membranes were stained with Ponceau-S solution (0.1 % Ponceau red dye (Fisher; BP103), 5% Acetic acid (Sigma;33209). Next, membranes were blocked for 1 h in 5% Milk (Coop;Milkpowder) in TBS-T (0.02 M Tris, 0.15 M NaCl, pH7.6 adjusted with HCl, 0.1% Tween20). Primary Antibody was added in appropriate dilutions in 5% Milk in TBS-T and incubated over night at 4°C. Membranes were washed three times for 10 min in TBS-T, before incubation with secondary antibody, diluted 1:10’000 in 5% Milk in TBS-T for 1 h at RT. Membranes were again washed three times for 10 min in TBS-T. Membranes were imaged on an Odyssey DLx infrared imager (LICORbio). Western blot quantification was done using Empiria studio software (LICORbio). Antibody list provided in Supplementary Table 1.

### RNA-seq analysis

Sequencing reads were trimmed using cutadapt^64^ v3.5 (removal of truseq adapter sequences, polyA sequences, and low-quality reads). Trimmed reads were aligned to the mouse genome (mm10) using STAR^65^ v2.7.5 (STARoptions, --readFilesCommand zcat --runThreadN 8 --outFilterMultimapNmax 20 --alignSJoverhangMin 8 --alignSJDBoverhangMin 1 --outFilterMismatchNmax 999 --outFilterMismatchNoverLmax 0.6 --alignIntronMin 20 --alignIntronMax 1000000 --alignMatesGapMax 1000000). Gene count matrices were obtained from the aligned reads using featureCounts^66^ v2.0.3 (-t exon -g gene_id -O -T 4 -s 1 -a GRCm38.102). Differential gene expression analysis between conditions was determined using the R package DESeq2^67^ after pre-filtering genes with low counts (rowSums > 10), with significance cut-offs set at FDR < 0.05 and fold change > 1.5 in either direction. Output data were plotted with ggplot2.

### Statistical analysis

The statistical tests performed are disclosed in the corresponding Figure legends. Statistical tests were performed using Prism and R.

## Data and materials availability

The next-generation sequencing data generated in this study will be deposited on the GEO server. The mass spectrometry proteomics data generated in this study will be deposited in the ProteomeXchange Consortium via the PRIDE partner repository database. All analyses were performed using previously published or developed tools, as indicated in Methods. Requests for materials should be addressed to S.B.

## Acknowledgments

We would like to thank F. Steiner, M. Stoeber, J. Kirkland, E. Bru, and N. Thoma for valuable discussions, advice and protocols. We kindly acknowledge members of the following UNIGE Faculty of Medicine platforms: Animal facility, Biomaging platform, Proteomics platform, iGE3 genomics platform and RE.A.D.S platform for helping to run experiments as well as members of High Performance Computing (HPC) team at UNIGE for their support. We thank our colleagues and group members in the Department of Genetic Medicine and Development for critical comments on the project and manuscript. This work was supported by the Swiss National Science Foundation grant PCEFP3_194305 (awarded to S.M.G.B.), the Olga Mayenfisch foundation (Zurich, Switzerland) and a Swiss Government Excellence Scholarship (awarded to Y. S.).

## Author contributions

Y.S. designed and conducted experiments, analyzed data, designed figures and co-wrote the manuscript. H.S. developed cell lines and analyzed data. L.A. performed KO experiments in ESCs and iNeurons. S.B. conceived the project, designed and conducted experiments, analyzed data and wrote the manuscript.

