## Supplementary Figures for "Redundancy masks functional specificity of SMARCD paralogs in neurodevelopment"

**Supplementary Figure 1.** Related to Figure 1.

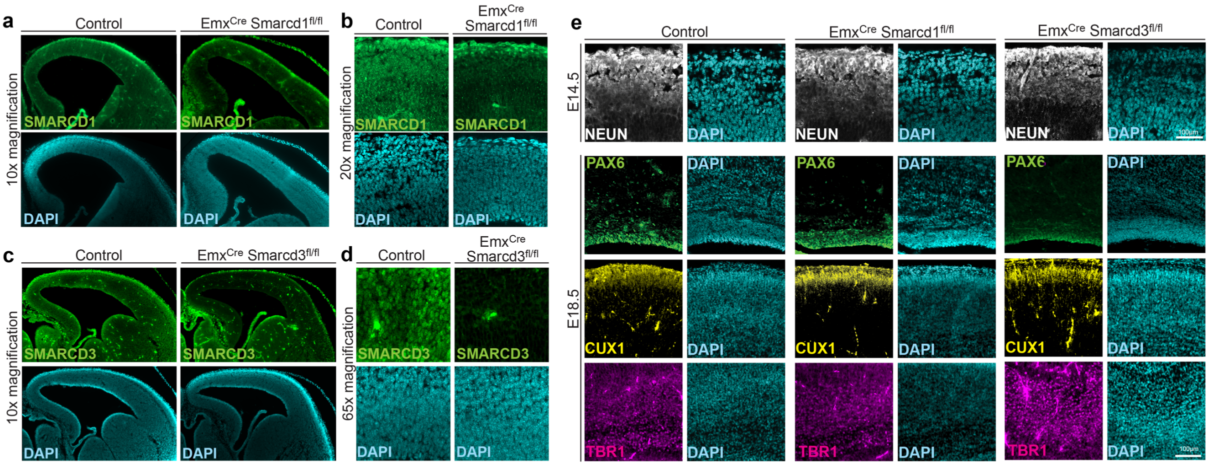


**a.** Representative images of coronal brain sections from control and *Smarcd1* cKO embryos at E14.5, showing Emx1Cre mediated forebrain specific deletion of *Smarcd1*. Sections were stained for SMARCD1. DAPI was used to visualize nuclei. Scale bars: 50 µm.
**b.** 20x magnification of SMARCD1 depletion in cortex of *Smarcd1* cKO embryos.
**c.** Representative images of coronal brain sections from control and *Smarcd3* cKO embryos at E14.5, showing Emx1Cre mediated forebrain specific deletion of *Smarcd3*. Sections were stained for SMARCD3. DAPI was used to visualize nuclei. Scale bars: 50 µm.
**d.** 65x magnification of SMARCD3 depletion in cortex of *Smarcd3* cKO embryos.
**e.** Representative images of coronal brain sections from control and *Smarcd1* cKO and *Smarcd3* cKO embryos at E14.5 and E18.5. Sections were stained for NEUN at E14.5, and for PAX6, CUX1, and TBR1 at E18.5. DAPI was used to visualize nuclei. Scale bars: 100 µm.

**Supplementary Figure 2.** Related to Figure 2.

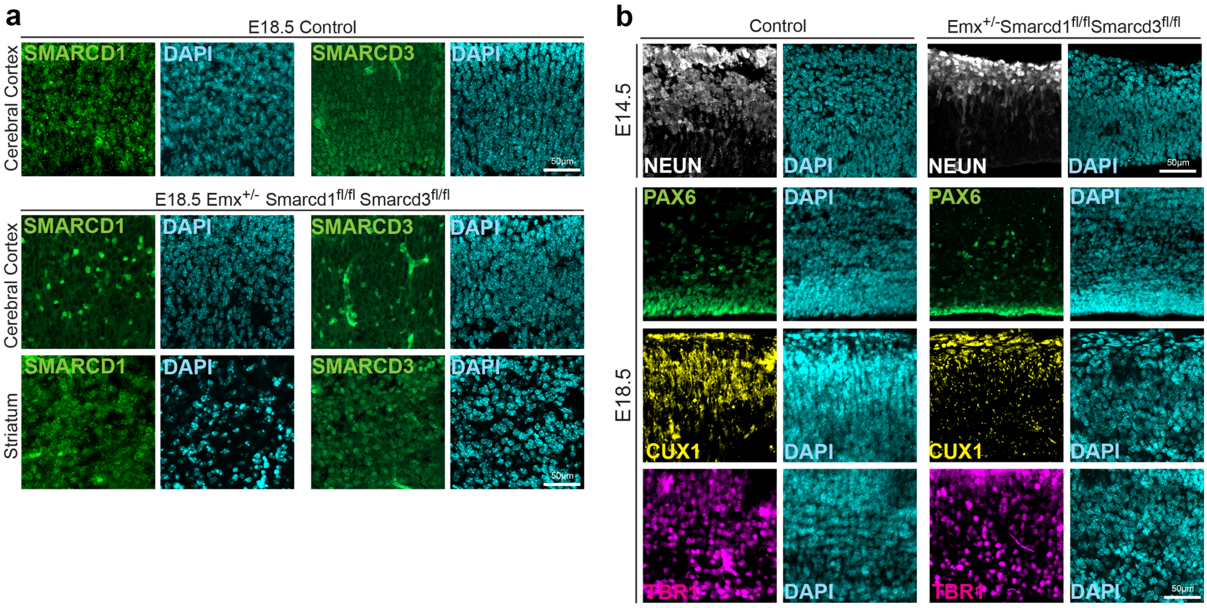


**a.** Representative images of coronal brain sections from control and *Smarcd1/3* double cKO embryos at E18.5, showing Emx1Cre mediated forebrain specific deletion of *Smarcd1* and *Smarcd3*. Sections were stained for SMARCD1 and SMARCD3. DAPI was used to visualize nuclei. Scale bars: 50 µm.

**b.** Representative images of coronal brain sections from control and *Smarcd1/3* double cKO embryos at E14.5 and E18.5. Sections were stained for NEUN at E14.5, and for PAX6, CUX1, and TBR1 at E18.5. DAPI was used to visualize nuclei. Scale bars: 50 µm.

**Supplementary Figure 3.** Related to Figure 3.

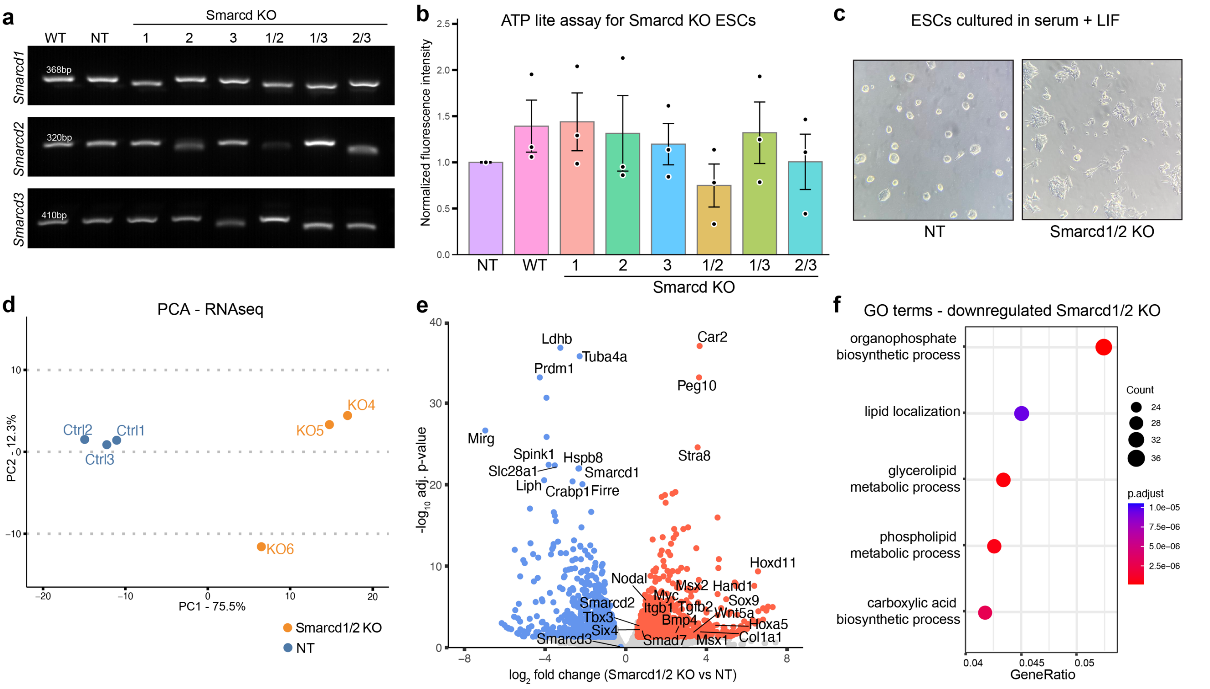


**a.** Genotyping PCRs of *Smarcd1/2/3* genes in *Smarcd* single and double knockout ESC lines. Two sgRNAs were used to target each allele, with homozygous deletion causing a shift in the PCR product as a result of the targeted deletion. Mutant allele PCR bands were analyzed by Sanger sequencing to confirm indels and KO.
**b.** ATPlite assay for *Smarcd* single and double KO and control ESCs. Plot shows fluorescence intensity normalized to NT condition. n=3. No significant differences $p > 0.05$ using a two-sided t-test.
**c.** Representative images of *Smarcd1/2* double knockout ESCs and NT controls, highlighting premature differentiation and loss of ESC-like morphology in KO cells.
**d.** Principal component analysis (PCA) of RNA-seq in *Smarcd1/2* double KO ESCs and NT controls. PCA was performed on variance-stabilized RNA-seq counts to assess global transcriptomic variation in ESCs. Each point represents a biological replicate (n = 3 per condition).
**e.** Volcano plots depict differentially expressed genes from RNA-seq in *Smarcd1/2* knockout ESCs. Each point represents a gene, with log2-transformed fold change on the x-axis and –log10-transformed adjusted P-value on the y-axis. Genes meeting significance thresholds (adjusted p-value < 0.05 and | FC | > 1.5) are shown in red (upregulated) or blue (downregulated); non-significant genes are shown in grey. Differential expression analysis was performed using DESeq2, n = 3.
**f.** GO term analysis of downregulated genes in *Smarcd1/2* knockout ESCs. Point size corresponds to gene count and colour represents adjusted p-values. The x-axis shows the gene ratio.

**Supplementary Figure 4.** Related to Figure 4.

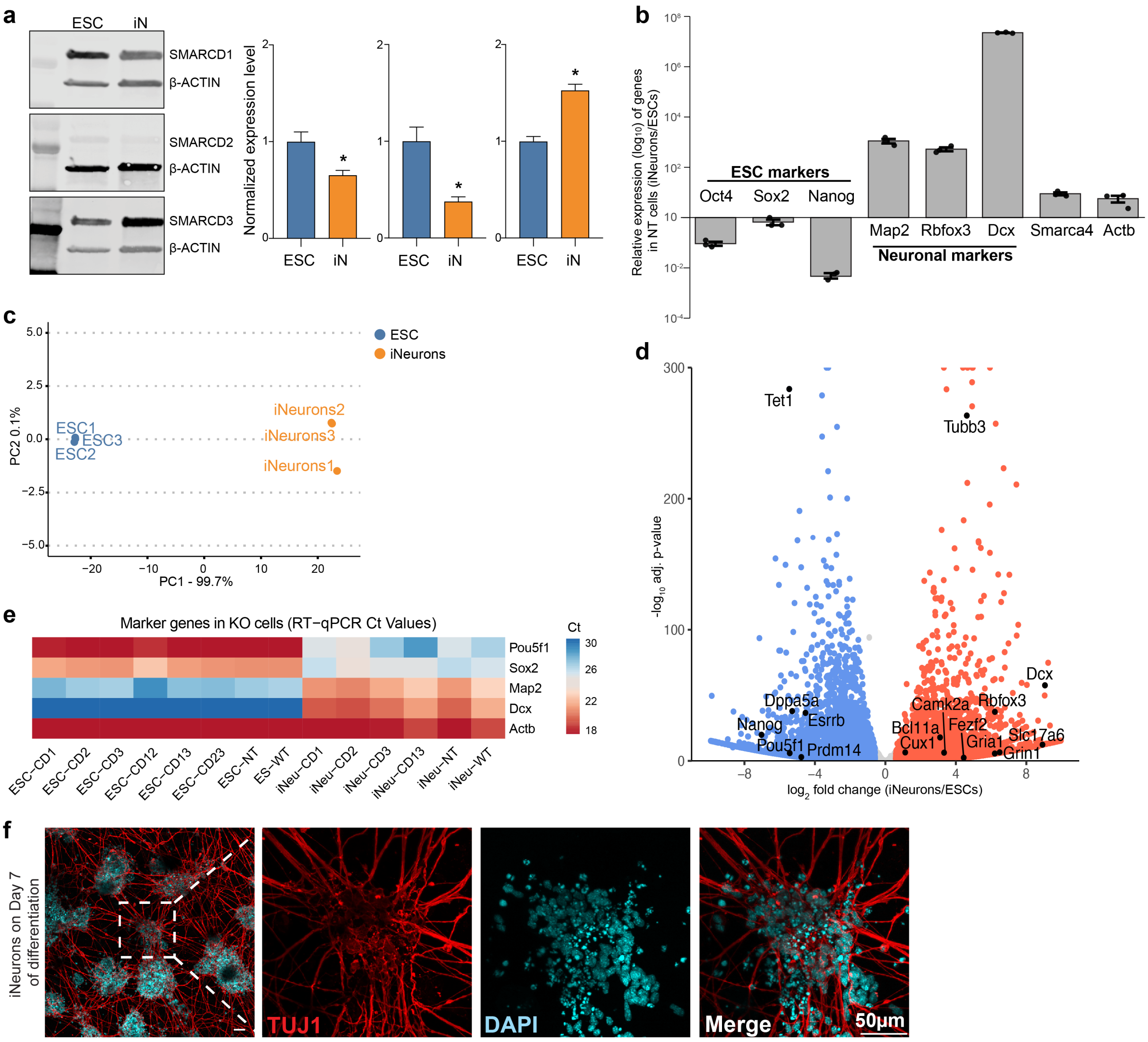


**a.** Western blot showing endogenous protein expression levels of SMARCD1, SMARCD2, and SMARCD3 in wild type ESCs and in vitro differentiated iNeurons. Quantification shows dynamic changes in SMARCD paralog levels across both cell types. β-actin was used as a loading control. n=3. P-value < 0.05 using a two-sided t-test.
**b.** RT-qPCR analysis of pluripotency (Oct4, Sox2, and Nanog), neuronal (Map2, NeuN, and Dcx) markers and SWI/SNF subunits (Smarca4 and Actb) showing relative expression in NT control iNeurons compared to NT control ESCs. y-axis shows the relative expression in log10 scale. n=3. ± SEM.
**d.** PCA was performed on variance-stabilized RNA-seq counts to assess global transcriptomic variation in NT ESCs and iNeurons. n = 3.
**d.** Volcano plots depict differentially expressed genes from RNA-seq in NT ESCs and iNeurons. Each point represents a gene, with log2-transformed fold change on the x-axis and –log10-transformed adjusted P-value on the y-axis. Genes meeting significance thresholds (adjusted p-value < 0.05 and | FC | > 1.5) are shown in red (upregulated) or blue (downregulated); non-significant genes are shown in grey. n=3.
**e.** Heatmap of RT-qPCR data showing differentially expressed genes in all *Smarcd* KO and control ESCs and iNeurons. The Ct values are depicted; low Ct values indicate highly expressed genes (red) and high Ct values indicate lowly expressed genes (blue). Actb Ct values serve as controls.
**f.** Representative images of iNeurons on day 7 of differentiation. Neurons were stained for TUJ1. DAPI was used to visualize nuclei. Scale bars: 50 µm

**Supplementary Figure 5.** Related to Figure 5.

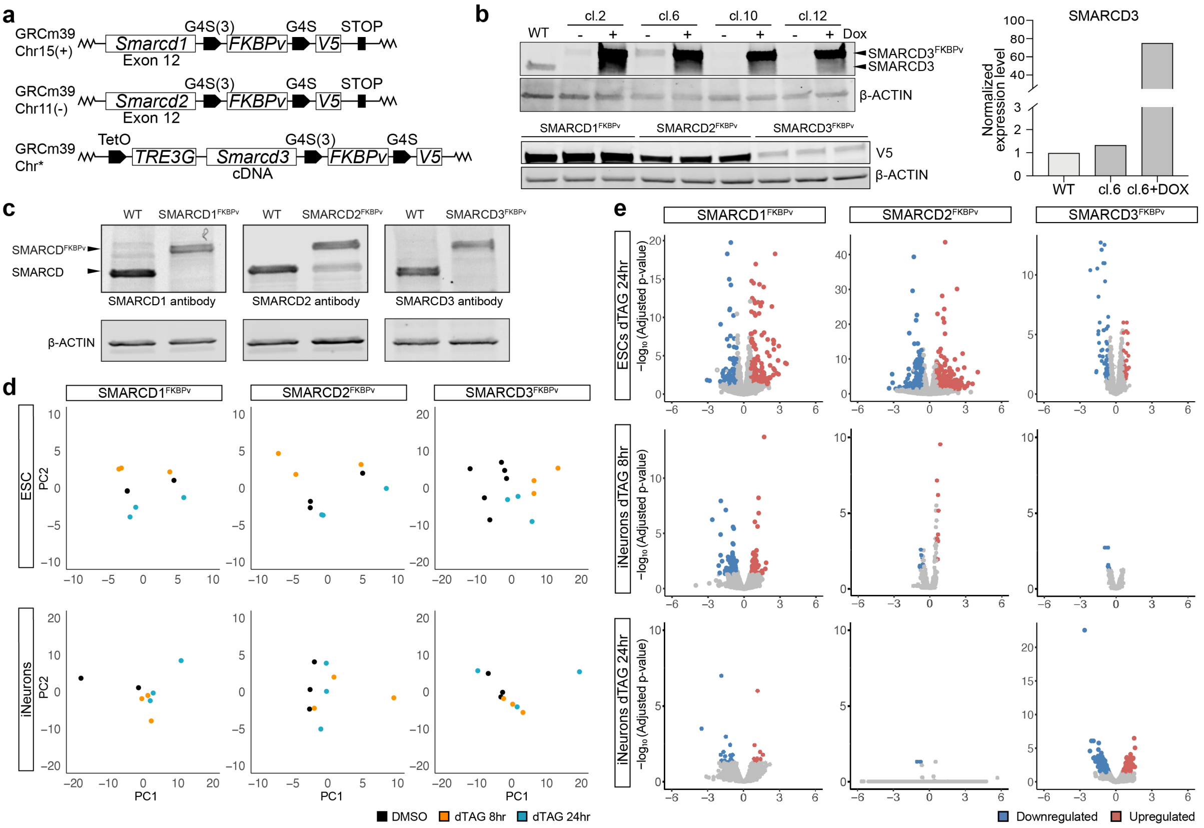


**a.** Schematic of the CRISPR/Cas9 knock-in strategy used to endogenously tag SMARCD1 and SMARCD2 at the C-terminus. An FKBPv degron domain followed by a (G4S)3 linker and a codon-optimized V5 epitope was inserted in-frame immediately upstream of the stop codon in exon 12, using homology-directed repair with plasmid-based donor constructs and locus targeting sgRNAs. For SMARCD3, repeated CRISPR knock-in attempts failed. Instead, we achieved biallelic disruption and generation of a *Smarcd3* knockout line. To restore *Smarcd3* expression, a doxycycline-inducible PiggyBac transgene encoding C-terminally FKBPv-V5 tagged SMARCD3 was stably integrated under the control of a TRE3G promoter. Notably, the transgene exhibited leaky expression at near-endogenous levels even in the absence of doxycycline, compensating for the knockout and enabling dTAG-mediated degradation.

**b.** Western blot shows SMARCD3$^{FKBPV}$ protein levels in selected clones (clones 2, 6, 10 and 12) and WT cells, with or without doxycycline treatment for 24 h. Clone 6 exhibits low-level leaky expression of SMARCD3$^{FKBPV}$ in the absence of doxycycline, whereas doxycycline induction results in robust overexpression of the fusion protein. Bar graph (right) shows quantification of SMARCD3 protein levels in WT and clone 6 SMARCD3$^{FKBPv}$ ESCs. β-actin was used as a loading control. Lower panel, western blot shows relative levels of SMARCD$^{FKBPv}$ fusion proteins in the three ESC degron lines using an antibody against the V5-tag.

**c.** Western blot analysis shows expression levels of SMARCD$^{FKBPv}$ fusion proteins relative to the untagged version of these proteins in WT ESCs. Note: the SMARCD2 antibody detects a non-specific band at the same height as the untagged SMARCD2 protein. We validated by genotyping PCR and Sanger sequencing that all SMARCD lines have homozygous in-frame knock-ins of the FKBPv-V5 tags.

**d.** PCA was performed on variance-stabilized RNA-seq counts to assess global transcriptomic variation in ESCs and neurons across three cell lines (SMARCD1$^{FKBPv}$, SMARCD2$^{FKBPv}$, and SMARCD3$^{FKBPv}$) under three treatment conditions: DMSO control, dTAG-13 treatment for 8 hours, and dTAG-13 treatment for 24 hours. Each point represents a biological replicate. One DMSO control replicate from the SMARCD1 neuron condition was identified as a strong outlier and excluded from the analysis. For the SMARCD3 ESC condition, DMSO controls include six replicates, as the 8-hour and 24-hour dTAG-13 treatments have different control samples. PCA was conducted using DESeq2-transformed counts, $n \geq 3$.

**e.** Volcano plots depict differentially expressed genes from RNA-seq in three SMARCD$^{FKBPv}$ cell lines in ESCs and iNeurons after 8 and 24 hours of dTAG-13 induced degradation. Each point represents a gene, plotted by log2 fold change on the x-axis and log10 adjusted p-value on the y-axis. Genes meeting significance thresholds (adjusted p-value < 0.05 and | FC | > 1.5) are shown in red (upregulated) or blue (downregulated), with non-significant genes in grey. $n \geq 3$.

**Supplementary Figure 6.** Related to Figure 6.

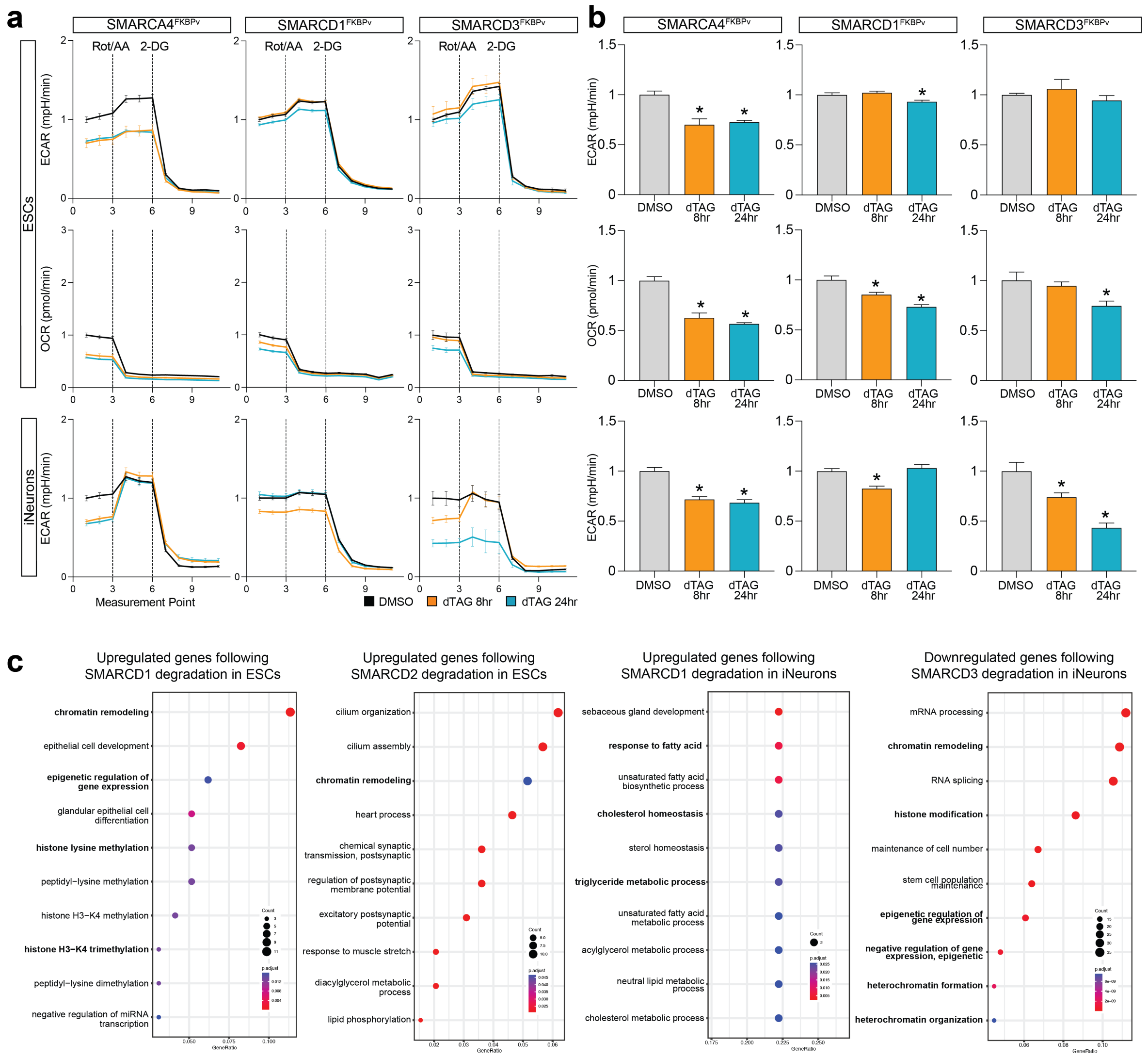


**a.** Glycolytic rate was measured at 11 sequential time points in SMARCD1$^{FKBPv}$, SMARCD2$^{FKBPv}$, and SMARCD3$^{FKBPv}$ cell lines in both ESC and iNeurons. The x-axis shows measurements 1–11, with tick marks 3, 6, and 9, corresponding to 14.2, 33.5, and 52.9 min, respectively. Cells were treated with -13-13 molecules for either 8 h (orange) or 24 h (blue), and compared to DMSO-treated controls (24 h). ECAR and OCR were measured.
**b.** Bar plots showing OCR and ECAR as the average of the first three points prior to rotenone and antimycin A. Data represent mean ± SEM; n ≥ 4 biological replicates per condition. Statistical significance was assessed using two-way repeated measures ANOVA to evaluate the effect of dTAG-13 treatment at the basal level and are indicated by asterisks (p-value < 0.05).
**c.** Key GO term analysis of upregulated and downregulated genes in SMARCD$^{FKBPv}$ cells after 24 hours of dTAG-13 treatment. Point size reflects gene count and colour indicates adjusted p-values. The x-axis shows the gene ratio.

**Supplementary Figure 7.** Related to Figure 6.

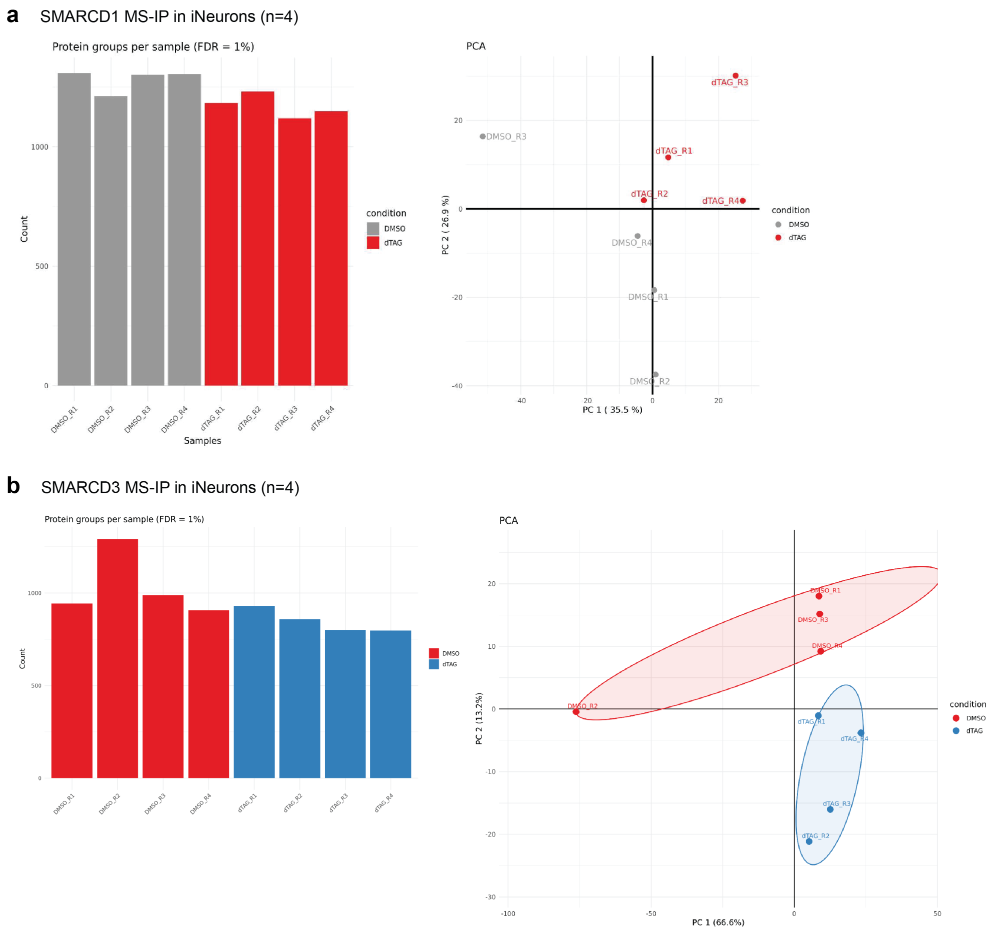


**a.** Immunoprecipitation mass spectrometry analysis of SMARCD1$^{FKBPv}$ interacting proteins in iNeurons. Bar plot showing number of identified proteins in each control and dTAG-13 treated replicate (left). PCA was performed on normalised protein levels (right). n=4.
**b.** Immunoprecipitation mass spectrometry analysis of SMARCD3$^{FKBPv}$ interacting proteins in iNeurons. Bar plot showing number of identified proteins in each control and dTAG-13 treated replicate (left). PCA was performed on normalised protein levels (right). n=4.

### Supplementary Tables.

**Supplementary Table 1.** List of primers and antibodies used in this study.
